# Differential maturation in vestibular neuronal groups related to developmental motor reorganization in amphibians

**DOI:** 10.64898/2026.05.12.724497

**Authors:** G. Barrios, A. Olechowski-Bessaguet, L. Cardoit, T. Fevrier, A. Wattignier, H. Tostivint, D. Cattaert, M. Thoby-Brisson, F. M. Lambert

## Abstract

Vestibular neurons are core elements of the pathways involved in vestibulo-motor functions, such as vestibulo-spinal and vestibulo-ocular reflexes. To meet behavioral needs, electrophysiological neuronal properties are adequately adapted to the sensory-motor computation sustaining these distinct vestibular reflexes. During frog metamorphosis, there is a complete reorganization of the posturo-locomotor system while the oculomotor system remains minimally changed, probably associated to so far unknown changes in vestibular neuronal properties. We used this unique model to investigate the central developmental mechanisms underlying such a reconfiguration of vestibular-associated behaviors. Central vestibular neurons exhibit two types of electrophysiological phenotypes: tonic neurons with a continuous discharge and phasic neurons with a transitory discharge mainly due to the activation of Kv1.1 channel. Electrophysiological recordings and Kv1.1 immunolabeling of vestibulospinal (VS) and vestibulo-ocular (VO) neurons at both larval and juvenile stages revealed that the majority of VS neurons exhibited a tonic discharge in larvae but a phasic discharge in juvenile, while VO neurons remained mainly tonic throughout development. Changes in phasic and tonic neurons proportions in VS population are partly explained by neurogenesis. But we provide evidences that an electrophysiological phenotype switch is a concomitant developmental mechanism participating in the maturation of these central vestibular neurons. All together our results showed that the maturation process in central vestibular neuronal groups is highly related to the metamorphosis-induced remodeling of vestibulo-motor functions they are involved in, with the ultimate purpose of ensuring an adequate adaptation of neuronal elements properties to the developmental changes of behavioral constrains.

## INTRODUCTION

Gaze and postural stabilizing behaviors are produced by vestibular sensory-motor neuronal networks with adequate molecular and cellular components required to generate the appropriate neural computation. Central vestibular neurons, also called 2^nd^ order vestibular neurons (2°VN), represent the neuronal “hub” where most of the sensory-motor transformations necessary to build ocular and postural reflexive tasks like the vestibulo-ocular (VOR) and vestibulospinal reflexes take place (Straka and Dieringer, 2004; Straka et al., 2005; Angelaki and Cullen, 2008). Common to all vertebrates, 2°VN receive vestibular sensory inputs but are involved in different functional motor pathways: vestibulo-ocular (VO) neurons project to oculomotor centers including extra-ocular motoneurons and participate to the VOR (Straka et al., 2001, 2002; Straka and Dieringer, 2004, Bianco et al., 2012), while vestibulospinal (VS) neurons project to spinal cord to produce vestibulospinal reflexes that balance trunk and limbs (Straka et al., 2001; Kasumacic et al., 2010, 2015, Lambert et al., 2013, 2016; Hamling et al., 2024; Zhu et al., 2024). If all 2°VN sub-populations share common basic features (Straka et al., 2005; Beraneck and Lambert, 2020), they probably exhibit more specific characteristics depending on the vestibulo-motor pathways they belong to. Some studies reported a correlation between firing properties in the one hand and neurotransmitter phenotype and/or ion channel, neurofilament and synaptic proteins on the other hand (Bagnall et al., 2007; Kolkman et al., 2011; Kodama et al., 2020). A few studies examined maturation of firing properties in vestibular sub-groups in a narrow developmental context like perinatal periods (Gamkrelidze et al., 1998 and 2000; Popratiloff et al., 2003; Dubois et al., 2022), but independently of the behavioral maturation and/or adaptation of vestibular reflexes. How much adaptive plasticity in membrane properties and discharge dynamic of 2°VN is functionally related to the maturation of vestibulo-ocular and vestibulospinal reflexes during the development remains poorly investigated.

The amphibian metamorphosis offers a unique and natural plasticity context to address this issue. Indeed, during development, the vestibulospinal reflexes and pathways undergo important modifications induced by the complete re-modeling of the posturo-locomotor system from larvae to froglet (Beyeler et al., 2008; Olechowski-Bessaguet et al., 2020; Barrios et al., 2024). In contrast, the conservation of the oculomotor system through metamorphosis does not provoke massive changes of ocular reflexive behavior (Dieringer and Precht, 1982; Dieringer, 1987; Horn et al., 1986; Lambert et al., 2008; Bacqué-Cazenave et al., 2022). Consequently, it appears relevant to investigate basic electrophysiological properties in identified vestibulospinal and vestibulo-ocular 2°VN in relation to this function-dependent developmental sensory-motor context. It has been described in the literature that frog 2°VN exhibit two distinct functional phenotypes according to their membrane properties and their discharge dynamic: 1) phasic neurons present a fast-transitory, adaptive, discharge of a few spikes and 2) tonic neurons elicit a continuous non-adaptive firing activity (Straka et al., 2004; Beraneck et al., 2007). Functionally, phasic and tonic neurons act as band pass-filter and low-pass neuronal filters, respectively, and participate to the signal encoding of the movement or the position. It is obvious that the switch during metamorphosis from the tail-based axial system in larvae to the tetrapod system in froglet changes the sensory-motor signal processing for position and movement encoding. Consequently, the following question raises: how much electrophysiological properties in vestibular neurons are modified from the larval life to the froglet stage depending on the pathway they are involved in? To address this issue, we combined patch-clamp recording, neuroanatomical approaches and computational modeling to investigate electrophysiological properties and developmental origin of identified VS and VO neurons at both larval and juvenile stages. Our results show a distinct maturation patterns between VS and VO neuronal populations, with the implication of different developmental mechanisms, that must be relevant according to the different degree of reorganization of either the posturo-locomotor or the oculomotor systems, respectively.

## MATERIALS AND METHODS

### Animal’s care

Experiments were performed at larval stages 50-53 (N=91), pro-metamorphosis stages 55-58 (N=9), climax stages 59-61 (N=22) and post-metamorphosis juvenile stage 65-66 (N=123) of the South African clawed toad *xenopus laevis*. Animal stages were identified according to external body criteria described in Nieuwkoop and Faber, 1956. Animals were maintained at 20–22°C in filtered water aquaria under a 12h: 12h light/dark cycle. All procedures were carried out in accordance with, and approved by, the local ethics committee (protocols #2016011518042273 APAFIS #3612).

### Tissue preparation

For all experimental procedures described below, animals were anesthetized using tricaine methanesulfonate (0.05% MS-222, Sigma-Aldrich) and were dissected out in a sylgard bottom petri dish with an ice-cold Ringer solution oxygenated with carbogen (95% O2 and 5% CO2) and containing (in mM): 93.5 NaCl, 3 KCl, 30 NaHCO3, 0.5 NaH2PO4, 11 glucose and pH adjusted to 7.4. Forebrain, viscera, otic capsules and cartilage covering the central nervous system were removed before any other treatment.

### Retrograde labelling of central vestibular neurons

Central vestibular neurons were retrogradely labeled after successive application of 3-4 small crystals of fluorescent dye dextran-coupled (tetra-methyl rhodamine 3kD (RDA) or Alexa fluor dextran 488 or 647 10kD, Life technologies) in a tiny incision performed in the projection site, as previously described (Straka et al., 2001; Lambert et al., 2013; Olechowski-Bessaguet et al., 2020). Vestibulospinal neurons were labelled by injecting the dye crystal to the ventro-medial surface of the rostral hemi-cord at the level of the first ipsilateral segment. Vestibulo-ocular neurons were labelled by injecting the dye crystal to the ventral-medial surface of the brain, at the hindbrain-midbrain boundary, where the oculomotor nerve gets out of the brain. Excess dye was washed out with cold Ringer solution. Thereafter, the brainstem-spinal cord preparation was incubated in circulating ringer solution maintained at 16-17°C, controlled electronically (Peltier, DAGAN corporation HCC-100A), for at least 3 hours to perform retrograde labelling of neuronal cell bodies in vestibular nuclei.

### Slice preparation and electrophysiological recordings

After the dye migration, the isolated *in vitro* “brainstem - spinal cord” preparation was partially included in a low-melting temperature Agar block (4%, Sigma Aldrich). The Agar block was oriented with the brainstem dorsal side up and placed in the vibratome cutting chamber (Leica VT 1000S) filled with an oxygenated sucrose solution at 4 °C (1.15mM NaH2PO4, 2mM KCl, 7mM MgCl2, 0.5mM CaCl2, 26mM NaHCO3, 11mM glucose, 205mM sucrose). The dorsal part of the brainstem was removed using a vibrating microslicer to expose vestibular nuclei. Preparations were transferred into a recording chamber under an upright microscope (Nikon Eclipse FN1) and continuously perfused with oxygenated Ringer solution, at 18-20°C. RDA+ vestibular neurons were identified by their dorso-lateral anatomical position in the brainstem throughout rhombomeres 3- 6 and rhombomeres 1-3 for vestibulospinal and vestibulo-ocular neurons, respectively. Neurons were visualized with a fluorescence system at a wavelength of 570nm (Dual optoled supply) with a 40X water-immersion objective.

Microelectrodes were pulled from borosilicate glass capillaries GC150F-10 (1.5 OD x 0.86x100 L mm) on a pipette puller (Sutter Instrument Co. Model P-2000). Whole-cell patch clamp recordings of 220 RDA+ vestibular neurons were performed under voltage and current clamp conditions using an Axoclamp 2A amplifier (Molecular Devices, Berkshire, UK). Traces were filtered at 10KHz using a digitizer interface (Digidata 1440; Molecular Devices) and acquired with the PClamp10 software. Patch pipettes resistance ranged around 5–8 MΩ when filled with the intracellular solution composed of: 115 mM K-gluconate, 2 mM MgCl2, 2 mM EGTA, 10 mM HEPES, 2 mM MgATP, 0.2 mM NaGTP. Biocytin (0.01%) or Neurobiotin (0.04%) was included in the pipette solution for intracellular filling and later detection. Series resistance was measured during all recording, in response to hyperpolarizing pulse of −10mV and was typically comprised between 12 and 20mΩ. Traces were analyzed offline using Clampfit (Molecular Devices)

Vestibular neurons were first recorded in current clamp condition and exposed to a series of depolarizing current step injections (from +50pA to 1nA, 500ms long, 25pA or 50pA increment) to identify their discharge dynamics. Input resistance was measured in response to a hyperpolarizing voltage step from −80 to −50mV. In control conditions, gap free recording was monitored during 5-10 minutes in order to ensure stable recording and to exclude cells in which resting membrane potential was too variable. Two or three consecutive recordings of various current or voltage clamp protocols were realized to characterize the neuronal phenotype. All electrophysiological traces were analyzed using Clampfit, Dataview and Origin 8.5.and were exported in Illustrator for figure making.

### Pharmacology

Bath application of dendrotoxin-k (DTX-k, 100nM, Alomone Labs) or 4-aminopyridine (4-AP, 10µM, Sigma) recognized as a specific Kv1.1 antagonist at low concentration, was performed to investigate the implication of Kv1.1-related I_D_ current in discharge dynamics of vestibular neurons. XE991 dihydrochloride (15µM, Alomone Labs) was perfused in the bath to test the implication of the Kv7.2/3 related I_M_ current in phasic 2°VN discharge. The involvement of SK Ca^2+^-dependent current (I_SK_) and BK Ca^2+^-dependent current (I_BK_) was conjointly tested with bath application of Apamin (100-200nM) and Iberiotoxin (50-100nM), respectively.

### BrdU incubation

Groups of tadpoles at larval stage 42, 48, 50, 54, 58 were incubated in a solution of 10mM BrdU (5’-bromo-2’-deoxyuridine, Sigma Aldrich, PULSE) tank water for 24h according to protocols previously published (Auger et al., 2012; Locker and Perron, 2019). After the PULSE, animals were breaded normally in our animal facility until stage 65 (CHASE) where vestibulospinal or vestibulo-ocular neurons were labelled as described above in combination with immunolabeling of Kv1.1 (see below) and BrdU detection.

### Immunocytochemistry

After patch clamp recordings, brainstem preparations containing biocytin+ and RDA+ neurons were first fixed overnight with 4% paraformaldehyde (PFA) at 4°C, then transferred to a PBS azide solution until performing a Kv1.1 immunolabeling as described below. To assess the proportion of Kv1.1+ neurons on the entire vestibulospinal or vestibulo-ocular populations fluorescent dye injections were performed the same way than described above. After tracer migration brainstem preparations were fixed 2 hours in 4% PFA at room temperature then transferred in a 20% sucrose solution made in 0.1% PBS overnight at 4°C. Consecutive to the cryoprotection in the sucrose solution the tissue samples were embedded in tissue tek (VWR-Chemicals) frozen at −45°C in isopentane and stored at −80°C. Then 25 µm cross-sections were obtained using a cryostat CM3050 (Leica, Leica France) and carefully placed on slides. After several rinsing in phosphate-buffered saline (PBS) 0.1%, sections were exposed to a blocking solution of PBS containing 1% bovine serum animal and 0.3% Triton X-100 for 90 min at room temperature to avoid unspecific labelling. Samples were then incubated overnight at 4°C with the rabbit primary antibody anti-Kv1.1 subunit (1/200, APC-009, Alomone labs). After washing with PBS, sections were incubated with the secondary antibody (1/500, Alexa fluor 488 goat anti rabbit) for 90 min in the dark at room temperature. After additional washes, cross-sections were mounted in a medium containing 74.9% glycerol, 25% Coon’s solution (0.1 M NaCl and 0.01 M diethylbarbiturate sodium in PBS) and 0.1% paraphenylenediamide. Conjointly with the Kv1.1 immunolabeling, isolated preparations were incubated using Alexa Fluor 647 streptavidin to reveal RDA+ vestibular neurons filled with biocytin during the patch-clamp recording session. BrdU immunolabeling was performed the same way than the Kv1.1 immunolabeling described above. Images were acquired using a confocal microscope (Zeiss Ax10 Imager M2) and processed by ImageJ FIJI software and Adobe Photoshop CS5.

### ISH Probe synthesis

Complementary DNA fragments encoding for *X. laevis* NeuroD1 (NM_001092127), Kv1.1 (NM_001085777) and kv7.4 (XM_041582171) were amplified by RT-PCR (for primer sequences see Table 1) from adult *X. laevis* brain and spinal cord RACE-ready cDNA as previously described (Bougerol et al., 2015; Alejevski et al., 2021), subcloned into pGEM T-Easy vector (Promega) and sequenced for confirmation. Purified cDNA fragments (854 bp for *neurod1*, 883 bp for *kv1.1* and 820 bp for *kv7.4*) were used to synthesize digoxigenin (DIG)-labelled sense and anti-sense cRNA probes *in vitro* with the RNA labelling Mix (Sigma Aldrich) following the manufacturers’ instructions.

**Table 1:**
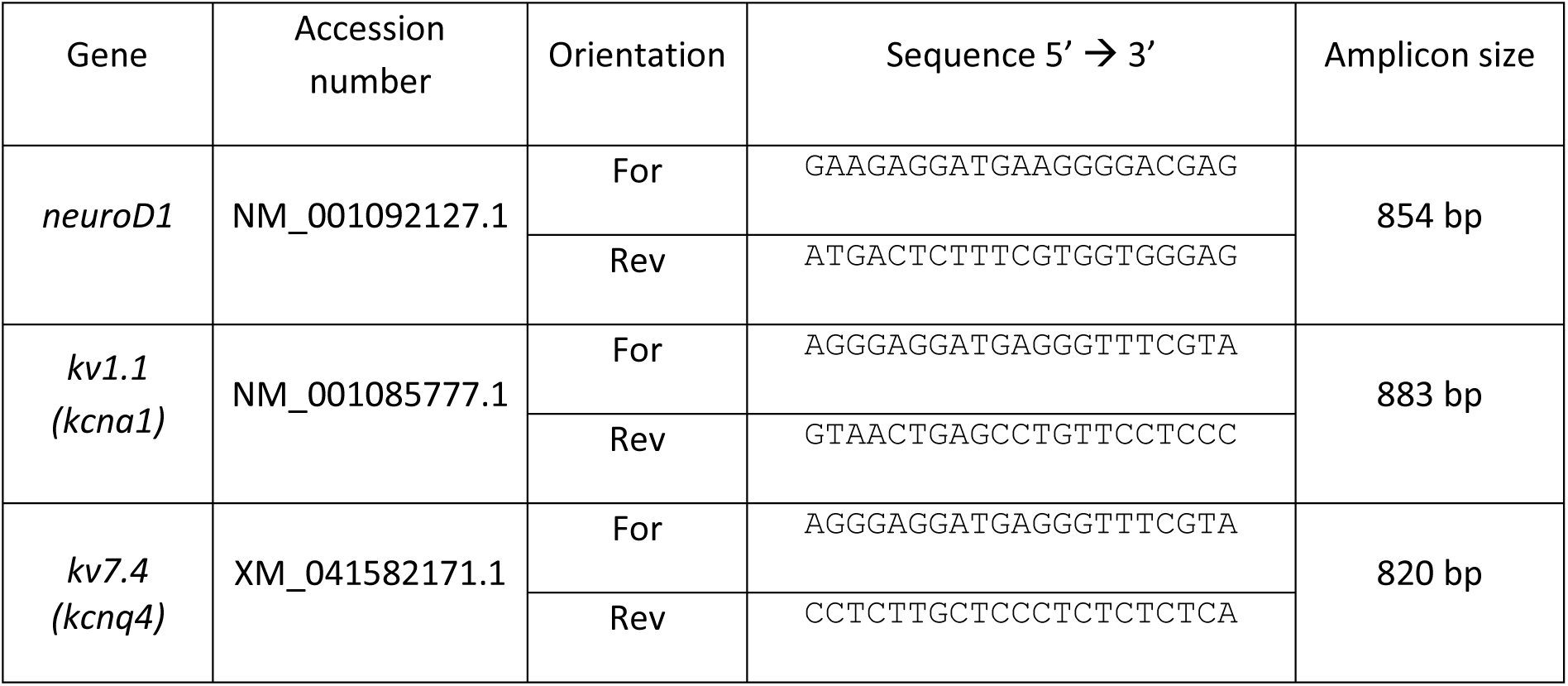
Primer Sequences for in-situ hybridization probe synthesis.

### In situ hybridization

15 µm brainstem cross-sections were obtained the same way than described above in RNAse free conditions. The *in situ* hybridization (ISH) protocol was adapted from previous studies (Bougerol et al., 2015; Lambert et al., 2018). In brief, after several rinses in PBS – 0.01 % Tween 20 (PBS-Tw) cross-sections were incubated in a prehybridization buffer (5 X SSC, 50 µg/ml heparin, 0.5 mg/ml yeast tRNA, 0.1% Tween-20, 9.2 mM citric acid pH 6.0) during 90 min at 64°c. After the prehybridization step, cross-sections were incubated overnight at 64°C in the same solution containing the heat-denaturized NeuroD or Kv1.1 or Kv7.4 riboprobe. After hybridization, sections were rinsed 3 times in 2 X SCC then one time in 0.1 X SCC for 60 min at 64°C. Two final washes were performed in a maleic acid buffer (100 mM maleic acid, pH 7.2, 150 mM NaCl, 0.1 % Tween-20) at room temperature. Sections were then incubated in a blocking solution [2.5% Blocking reagent (Roche), 5% Sheep serum in maleic acid buffer] at room temperature for 60 min before adding the primary antibody anti-fluo-POD solution (1/150 in the blocking solution) overnight at 4°C. After several rinses in PBS-Tw at room temperature, labeled probes were revealed by the Tyramide Signal Amplification method as previously described (Quan et al., 2015) using a home-made FITC- (Fischer scientific) conjugated tyramide. After extensive washes in PBS-Tw (at least five times for 20 min each), sections were mounted (Fluoromount mounting medium, Invitrogen) or underwent an immunolabeling protocol as described above. The specificity of the hybridization procedure was verified by incubating sections with the sense riboprobes with which only background signals typical of this type of chemical reaction could be observed (data not shown).

### Confocal image acquisition and image analysis

All neuroanatomical preparations with fluorescent signals from either immune- or ISH-labelings were acquired using a confocal microscope (Zeiss Ax10 Imager M2) equipped with 405, 488, 543 and 633 nm laser lines. Multi-image stacks of images with a 1-3µm Z-step interval were obtained with a 20x/0.75 oil objective and with a 0.3-0.5µm Z-step interval with a 60x/1.4 oil objective. Confocal images processed with artificial fluorescent colors in the FIJI software environment and Adobe Photoshop CS5. The counting of 2°VN populations Kv1.1 and/or BrdU immune positive or negative was performed by hand through multi-images stacks by the same person to avoid counting bias. The level of the NeuroD expression in brainstem cross-sections was evaluated by calculating the ratio between the area of the NeuroD fluorescent signal and the total cross-section area.

### Statistics

Average data were expressed as mean ± SD unless mentioned otherwise. Assumption of normality was checked with Shapiro-Wilk test and sample variances compared with F-Test. All statistical tests are indicated in figures legends. The level of significance was * = p<0.05, ** = p<0.01, *** = p<0.01.

### Computational modeling

A basic computational model of 2°VN was created with the NEURON 8.2.4 program (Hines and Carnevale, 1997), including a soma (diameter = 20µm), a dendrite (diameter = 2µm; length = 500 µm), a passive element (diameter = 2µm; length = 50 µm) leading through an axonal initial segment (AIS, diameter = 2µm; length = 90 µm) to the axon (diameter = 2µm; length = 500 µm). The number of segments was prescribed an odd number that was calculated according to the d-lambda rule (Carnevale and Hines, 2006; Hines and Carnevale, 2001). A passive leakage current (I_leak_) was inserted in all sections. I_leak_ was simulated as followed [I_leak_ = (E_leak_ – E) × G_leak_, with E_leak_ = −73 mV and G_leak_ = 1/R_m_], where *gleak* (in mS/cm²) is the constant membrane conductance associated with non-gated “leak” channels and *E_leak_* is the effective reversal potential for the leak current, i.e. the membrane voltage at which the net leak current is zero. R_m_, the specific membrane resistance, was set to R_m_=10 MΩ in axonal sections and R_m_=20 MΩ in all other sections. In addition, calcium dynamics were added in the soma to reproduce calcium accumulation, diffusion and pumping (Destexhe et al., 1993). Intracellular calcium dynamics were adapted from Destexhe et al. (1993) and set with the following parameters: depth = 0.0029 mm, taur = 1e10 ms, CainF = 0.0002 mM, Kt = 0.0025 mM/ms, Kd = 0.008 mM, Cai = 2 × 10−4 mM and Cao = 2 mM. The following ion channels were inserted in the membrane of specific cellular sections according to previous studies: Na, Kv3.3 (Kht) and KV1.1 channels from Watanabe et al. (2017); KV7.2/3 from Shah et al. (2008); CaN from Benison et al. (2001); BKCa from Mangde&Manchanda (2018); SKCa from Mahapatra, et al (2018). Maximum conductance is expressed in in S/cm² in Supplemental Table 1.

## RESULTS

### Electrophysiological phenotypes of vestibular neurons

Vestibulospinal (VS) neurons, retrogradely labeled with a Dextran dye injection in the rostral hemi-cord, were located dorso-laterally in the brainstem (Fig. 1A). According to previous descriptions (Straka et al., 2001; Lambert et al., 2013; Olechowski-Bessaguet et al., 2020; Barrios et al., 2024), an ipsilateral population extended from the rhombomere 3-4 boundary to the rhombomere 6 forming the Lateral Vestibulospinal Tract (LVST, Fig. 1A), and a contralateral population was restricted to rhombomeres 5-6 forming the Tangential nucleus (TAN, Fig. 1A). In the present study we mainly focused our investigations on VS neurons from the LVST. However, neurons from the TAN were occasionally recorded and no obvious difference with LVST neurons was observed. Vestibulo-ocular (VO) neurons were retrogradely labeled with a unilateral dye injection in the oculomotor nucleus, at the midbrain-hindbrain boundary. Consistently with previous descriptions (Straka et al., 2001), VO neurons extended bilaterally in a dorso-lateral position but mainly neurons of the ipsilateral population, from rhombomere 1 to 4, were recorded in this study (Fig. 1A). Again, a few contralateral VO neurons were sporadically recorded without showing any particular differences compared to the ipsilateral group. The previous characterization of vestibular central neurons established in adult frog (Straka et al., 2004; Beraneck et al., 2007), was used to classify 2°VN based on their highly distinguishable discharge pattern. For instance, phasic neurons exhibited a transitory short adaptative discharge in response to current step injections, with a maximum of 5 spikes generated at the onset of the stimulation (Fig. 1B_1-2_). Inversely, tonic neurons fired continuously until the end of the current step (Fig. 1C_1-2_). These two phenotypes were found in both VS and VO neuronal populations and at both larval and juvenile stages (Fig. 1B-C).

**Figure 1.**
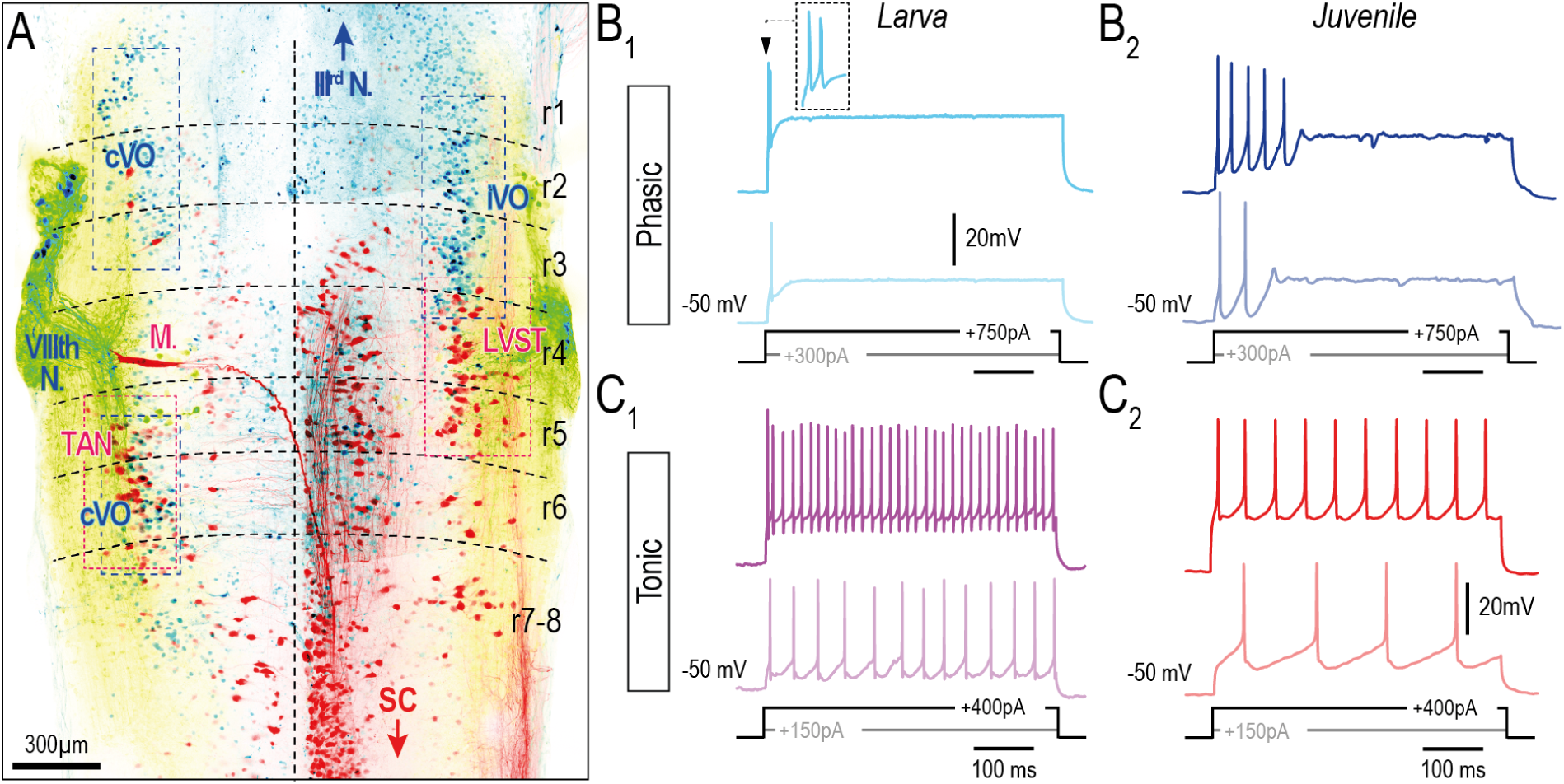
Electrophysiological phenotypes in vestibulospinal and vestibulo-ocular 2°VN in Xenopus frog. **A.** Confocal stack image of a whole-mount larval brainstem showing ipsi- (LVST) and contralateral (TAN) vestibulospinal (in red), ipsi-(iVO) and contralateral vestibulo-ocular (cVO; in blue) neurons and vestibular afferences (green) retrogradely labeled from the right hemi-spinal cord (SC, indicated by a red arrowhead), from the right oculomotor nucleus (III_rd_ N. indicated by a blue arrowhead) and bilateral VIIIth nerves (VIIIth N.). Dashed lines represent the midline and the rhombomere (r) boundaries. **B-C.** Examples of discharge dynamics in larval (B_1_-C_1_) and juvenile (B_2_-C_2_) phasic (B_1_-B_2_) and tonic (C_1_-C_2_) 2°VN in response to increasing positive current step injections during patch-clamp recording. M. = Mauthner cell.

**Figure 2.**
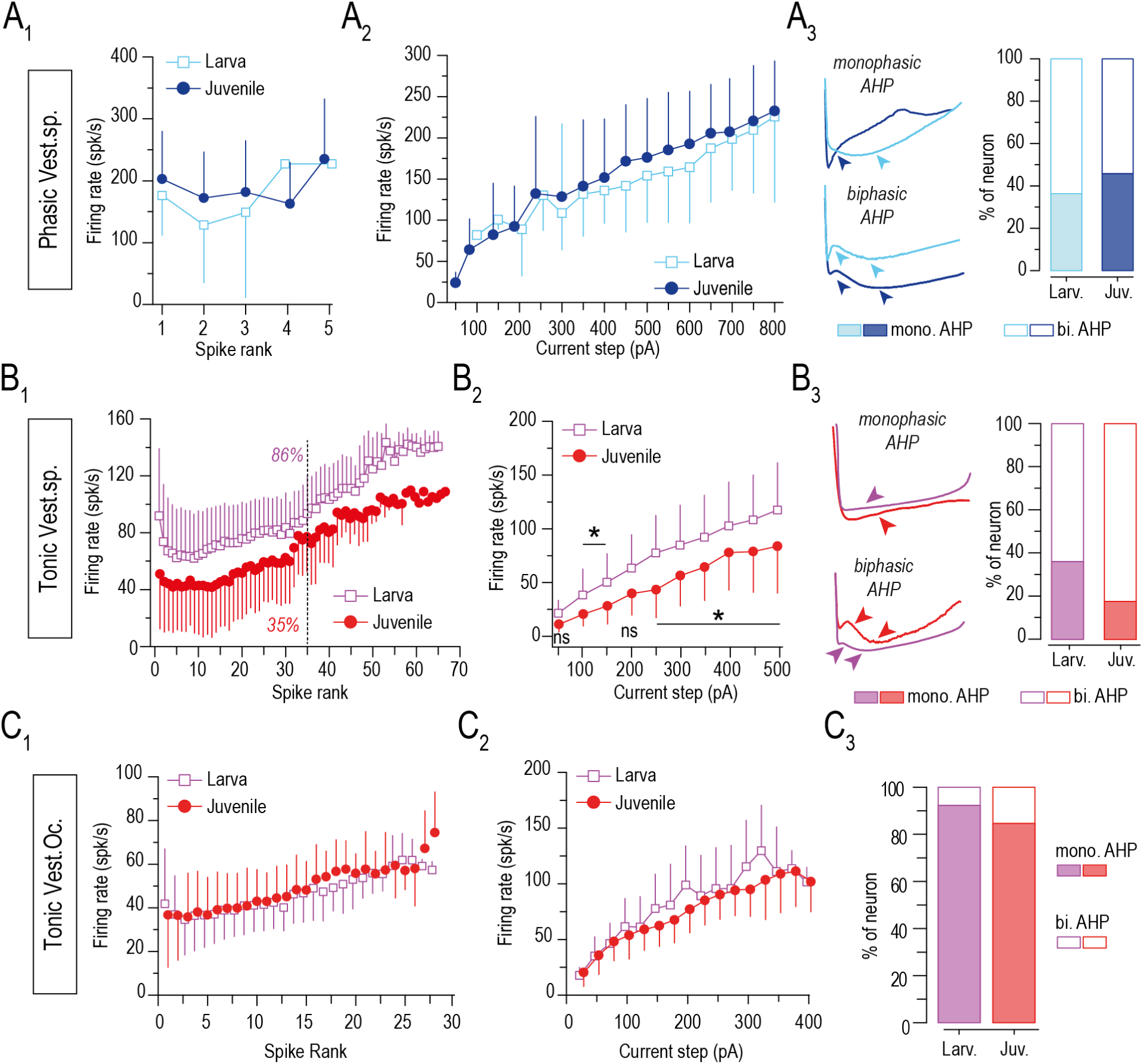
Firing properties of phasic and tonic vestibulospinal and vestibulo-ocular neurons in larval and juvenile xenopus. **A.** Averaged firing rate (±SD; in spike/s) plotted against the spike rank (A_1_, the position of the spike in the spike train) or against the injected current step (A_2_) and proportion of mono- and biphasic (bi.) after-hyperpolarization (A_3_; AHP; see examples of mono-and biphasic AHP at both stages on the left) in phasic vestibulospinal (vest.sp.) at both larval (cyan; larv.) and juvenile (dark blue; juv.) stages. **B.** Average firing rate (±SD; in spike/s) plotted against the spike rank (B_1_) or against the injected current step (B_2_) and proportion of mono- and biphasic AHP (B_3_) in tonic vestibulospinal 2°VN at both larval (purple) and juvenile (red) stages. Percentages in B_1_ indicate the proportion of tonic neurons firing less than 35 spikes, half of the maximum number of spikes elicited by tonic neurons. **C.** Average firing rate (±SD; in spike/s) plotted against the spike rank (C_1_) or against the injected current step (B_2_) and proportion of mono- and biphasic AHP (B_3_) in tonic vestibulo-ocular (vest.oc.) 2°VN at both larval (purple) and juvenile (red) stages. Statistical tests: Assumption of normality was checked with Shapiro-Wilk test and sample variances compared with F-Test. Firing rate of larval and juvenile neurons were compared with Student’s t-test or non-parametric equivalent Mann-Whitney U-test for each spike rank or current step (* = p<0.05); ns: not significant.

### Developmental Changes in discharge patterns of vestibular neurons

Phasic VS neurons presented comparable discharge pattern in tadpoles (before metamorphosis) and froglets (after metamorphosis, Fig. 2A). At both stages, 15% to 20% of recorded phasic VS neurons fired only one spike. For neurons exhibiting more than one spike, the firing rate was relatively constant around 180 spikes/s independently of the spike rank (calculated at 500pA in Fig. 2A_1_). The firing rate increased linearly from 75 to 225 spikes/s in response to current step injections from 100 to 800pA (Fig. 2A_2_). Phasic VS neurons were able to fire spikes with either a monophasic or a biphasic after hyperpolarization (AHP, Fig. 2A_3_) with some variations in the AHP shape (see examples shown in Fig. 2A_3_). The proportion of neurons with a monophasic AHP was a bit less in larva (∼36%) than in juvenile (∼45%). Overall, discharge dynamics of phasic VS neurons did not demonstrate significant changes from tadpole to froglet.

At the opposite, discharge pattern of tonic VS neurons presented some significant changes from tadpoles to froglet (Fig. 2B_1-3_). Interestingly, 21.4% (12/56) of larval tonic VS neurons showed a spontaneous discharge at rest, relatively regular (ySup. Fig. 2A-B), with an instantaneous firing rate of 8.4 ± 4.3 spike/s and a mean firing rate of 8.4 ± 4.0 spike/s (Sup. Fig. 2C). In contrast, in juvenile no tonic VS neurons was able to fire at rest. Tonic VS neurons responded rarely to current step injections above 500 pA. The firing rate increased with the spike rank at both stage but was lower and exhibited less spikes in juvenile than in larvae (Fig. 2B_1_). 86% of tonic VS neurons in larvae fired more than 35 spikes; half the maximum of 70 spikes measured, whereas only 35% of tonic VS neurons reached this spike threshold in juvenile (measured at 300 pA in Fig. 2B_1_). Consequently, the firing rate was also lower in response to increasing current steps in juvenile than in larvae (Fig. 2B_2_). Like phasic VS neurons, tonic neurons fired spikes with either a mono- or a biphasic AHP at both developmental stages and with some shape variations (Fig. 2B_3_). The proportion of neurons with a monophasic AHP was higher in larvae (∼36%) than in juvenile (∼17% only; Fig. 2B_3_). Altogether, these results showed a significant slower discharge dynamic of tonic VS neurons after metamorphosis.

Inversely to VS neurons, tonic VO neurons demonstrated very similar electrophysiological properties in tadpole and juvenile (Fig. 2C_1-3_). 41% (16/39) of larval tonic VO neurons showed a relatively regular spontaneous discharge at rest (Sup. Fig. 2A-B) with an instantaneous firing rate of 7.2 ± 4.3 spike/s and a mean firing rate of 7.1 ± 3.7 spike/s (Sup. Fig. 2C). In juvenile, only two tonic VO neurons exhibited a spontaneous firing activity at rest (Firing rate = 14 ± 0.4 spike/s). In response to current injections, the firing rate increased linearly from ∼40 to ∼55 spikes/s with the spike rank (for a current step of 300pA; Fig. 2C_1_) and from ∼25 spikes/s to ∼100 spikes/s in response to increasing current steps series (Fig. 2C_2_) at both developmental stages. Inversely to VS neurons, a large majority of tonic VO neurons evoked spikes with a monophasic AHP at both stages (90% in larvae, 85% in juvenile; Fig. 2C_3_), also contrasting with previous data reported in adult frog (Straka et al., 2004). Strikingly, only very few VO neurons with a phasic discharge could be recorded, 5 out of 32 at juvenile stages and 1 out of 40 in larvae, thus preventing any further robust statistical analysis (see examples in Sup. Fig. 2D). Note that most of phasic VO neurons fired spikes with a monophasic AHP (Sup. Fig. 2E). Overall, the analysis of the discharge pattern in 2°VN revealed major developmental changes exclusively in VS neurons, that are involved in the vestibular descending control. This significant adaptation of firing properties in tonic VS neurons might be related to the remodeling of spinal networks responsible for the postural behavior of the animal during the metamorphosis and the subsequent necessity to adapt vestibulospinal pathways (Olechowski-Bessaguet et al., 2020; Barrios et al., 2024).

### Proportion of phasic and tonic neurons in vestibulospinal and vestibulo-ocular populations before and after metamorphosis

The proportion of recorded phasic and tonic neurons was quite different from larvae to juvenile in one hand and from VS to VO populations in another hand (Fig. 3A). The most important developmental change was observed in VS neuron where the proportion of phasic and tonic phenotypes was significantly remodeled during the metamorphosis. Indeed, the VS populations contained a majority of tonic neurons in larvae (∼80%; Fig. 3A_1_) but a majority of phasic neuron in juvenile (∼70%, Fig. 3A_1_), revealing an important switch in the respective proportion of both phenotypes. At climax stage (stage 59-61), where both larval axial and juvenile appendicular posturo-locomotor systems co-exist (Combes et al., 2004), 40% of recorded VS neurons exhibited a phasic phenotype, twice than in larvae, strongly suggesting that the reversion of the proportion between phasic and tonic phenotypes might occur progressively during the metamorphosis. Comparatively to VS neurons, VO neurons remained mainly tonic throughout development (∼98% at larval stage and ∼85% at juvenile stage; Fig. 3A_2_), thus presenting a completely different developmental pattern than the vestibulospinal population.

**Figure 3.**
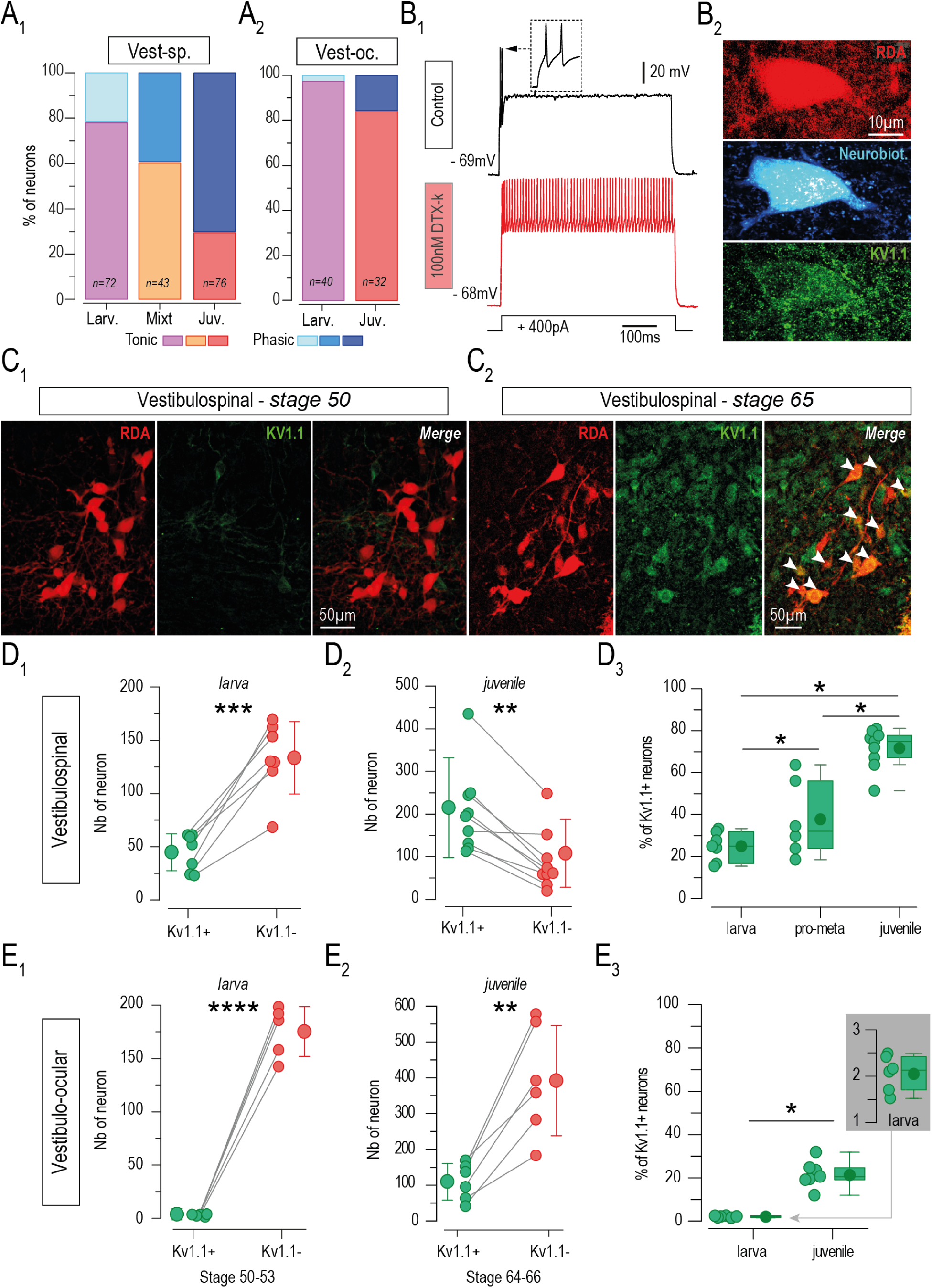
Proportions of phasic and tonic phenotypes in vestibulospinal and vestibulo-ocular neurons at larval, mixt and juvenile stages. **A.** Proportions of recorded phasic and tonic neurons in vestibulospinal (A1; vest-sp.) and vestibulo-ocular (A2; vest-oc) 2°VN in larvae (Larv.), mixt (climax stage) and juvenile (Juv.). **B.** Examples of discharge dynamics (B_1_) in a phasic 2°VN before (control, upper trace) and during 100nM DTX-k bath application (red lower trace) and confocal image (B_2_) of the same neuron retrogradely labeled with rhodamine (RDA, upper image in red), backfilled intracellularly with neurobiotin (Neurobiot.) during the path-clamp recording (middle image in cyan) and Kv1.1 immunolabeled (bottom image in green). **C.** Example of confocal images (cross-sections) showing vestibulospinal neurons retrogradely labeled with rhodamine dextran (RDA, in red) and immunodetection of Kv1.1 (green) at larval stage 50 (C_1_) and juvenile stage 65 (C_2_). White arrowheads in C_2_ indicated Kv1.1 positive VS neurons. **D.** Number of vestibulospinal neurons (D_1-2_) Kv1.1 negative (Kv1.1-, red dots) and positive (Kv1.1+, green dots) at larval (D_1_) and juvenile (D_2_) stages as well as proportions of Kv1.1+ VS neuron (D_3_) in larva (stage50-53) pro-metamorphosis (pro-meta; stage 54-58) and juvenile (stage 65) stages. **E.** Number of vestibulo-ocular neurons (E_1-2_) Kv1.1- (red dots) and Kv1.1+ (green dots) at larval (E_1_) and juvenile (E_2_) stages as well as proportions of Kv1.1+ VO neurons (E_3_) in larva (stage50-53) and juvenile (stage 65) stages. Statistical tests: Assumption of normality was checked with Shapiro-Wilk test and sample variances compared with F-Test. Numbers of Kv1.1+/- neurons (D_1-2_, E_1-2_) were compared with Student’s t-test or non-parametric equivalent Mann-Whitney U-test (*= p<0.05, **= p<0.01, ***=p<0.005, **** p<0.001). Percentages of Kv1.1+ neurons (D_3_, E_3_) were compared with Khi² test (*=p<0.05).

To confirm the proportions of the different phenotypes estimated from patch-clamp recordings we decided to perform an anatomical large-scale quantification over the entire LVST and the ipsilateral VO populations in the brainstem (see Fig. 1A). For this purpose, it was mandatory to select potential molecular markers specific of phasic and/or tonic phenotypes. For instance, the I_D_ conductance has been previously identified to be the main conductance involved in the transitory discharge of phasic vestibular neurons in adult frog (Beraneck et al., 2007), but also in chick embryo (Gamkrelidze et al., 1998 and 2000; Peusner et al., 1998). These past studies also demonstrated the predominant implication of the voltage-dependent potassium channel Kv1.1 in the I_D_ current, that could be further revealed by its sensitivity to dendrotoxin-k or to 4-aminopyridine at low concentration (DTX-k, 100 nM; 4-AP, 2-10µM; Peusner et al., 1998; Beraneck et al., 2007; Johnston et al., 2010). In addition, Kv1.1+ vestibular neurons can be immunolabeled in frog using a commercial anti-body against Kv1.1 (Beraneck et al., 2007; I Gusti Bagus et al., 2019). Therefore, these pharmacological and anatomical characterizations were used to confirm that Kv1.1 could be considered as a specific molecular marker to identify phasic KV1.1+ neurons in larval and juvenile preparations (Fig. 3B and Sup. Fig. 3). In phasic neurons, bath application of DTX-k (Fig. 3B_1_) or 4-AP (Sup. Fig. 3A-C) transformed the characteristic adaptive discharge of 1-5 spikes into a continuous firing activity comparable to that of tonic neurons. In the presence of these drugs, the number of spikes significantly increased above the expected maximum of 5 spikes (up to 50 spikes with DTX, Sup. Fig. 3B; up to 20 spikes with 4-AP; Sup. Fig. 3C), leading to the generation of a firing pattern with a discharge frequency at a range comparable to that of tonic neurons (Sup. Fig. 3B-C). Moreover, recorded phasic neurons were filled with neurobiotin to subsequently be simultaneously immunolabeled against Kv1.1. Phasic neurobiotin+ neurons recorded and treated with either DTX-k or 4-AP appeared to be Kv1.1+ (Fig. 3B_2_ and Sup. Fig. 3D). In contrast, and as expected, bath application of either DTX-k or 4-AP did not change significantly the discharge pattern of tonic neurons (Sup. Fig. 3E) and the number of spikes and the firing rate remained similar compared to control condition (Sup. Fig. 3F-G). Also, Kv1.1 immunolabeling revealed that none of the recorded tonic neurons were Kv1.1+ (Sup. Fig. 3H). Altogether, the combined pharmacological and immunolabeling characterizations demonstrated that Kv1.1 constituted a molecular marker specific of phasic neurons that could be used to distinguish the two vestibular phenotypes in VS and VO populations at a larger neuroanatomical scale.

Thus, Kv1.1 immunodetection was performed to identify phasic neurons, in the entire ipsilateral VS and VO populations, labeled by retrograde tracing at different developmental stages (Fig. 3C). The number of Kv1.1- VS neurons (red but not green; Fig. 3C), presumably tonic, was more than twice (∼134.1±34.0) the number of Kv1.1+ VS neurons (∼44.9±17.3, red and green; Fig. 3C) at larval stages 50-53 (n=7; Fig. 3C_1_ and D_1_). Inversely, the number of Kv1.1+ VS neurons (∼205.6.5±100.4) was about twice the number of Kv1.1- VS neurons (∼90.1±69.4) in juvenile (n=9; Fig. 3C_2_-D_2_). Consequently, the proportion of Kv1.1+ VS neurons was about 25% (24.9 ± 6.9) in larvae and about 70% (71.6±9.5) in juvenile (Fig. 3D_3_). Interestingly the proportion of Kv1.1+ VS neurons in pro-metamorphosis animals (stage 54-59) was around 40%, spreading from 20 to 65% (Fig. 3D_3_). A few VO neurons were Kv1.1+ at larval stage (3.6 ± 1.1; n=6; Fig. 3E_1_) representing a proportion of ∼2% (1.8 ± 0.5; Fig. 3E_3_) of VO neurons. The number of Kv1.1+ VO neurons was about ∼100 neurons in juvenile (106.3 ± 51.5; n=7; Fig. 3E_2_) corresponding to 20% of the total VO neurons at that stage (21.3% ± 6.6; Fig. 3E_3_). The histological quantification of Kv1.1+ and Kv1.1- neurons in the entire VS and VO population was therefore consistent with the proportions of phasic and tonic neurons identified from patch-clamp recordings. Indeed, VS population exhibited a majority of tonic Kv1.1- neurons (∼70-80%; Fig. 3A_1_ and 3D) in larvae but a majority of phasic Kv1.1+ neurons in juvenile (∼80%; Fig. 3A_1_ and 3D). The proportion of phasic Kv1.1+ VS neurons at the metamorphosis climax (∼40%; Fig. 3A_1_) and its great variability at the pro-metamorphosis (Fig. 3D_3_) suggests that the switch in the expression of phasic and tonic phenotypes in the VS group is a progressive plasticity mechanism occurring during a long period rather than a short developmental event restricted to a single larval stage. Inversely, VO neurons expressed mainly the tonic Kv1.1- phenotype before (Fig. 3E_1_) and after metamorphosis (Fig. 3E_2_), even if the proportion of phasic VO neurons increased a bit in juvenile (21.3% ± 6.6; Fig. 3E_3_). Altogether, these results demonstrated clearly that the expression of phasic and tonic phenotypes changed differentially in vestibular neuronal populations during frog metamorphosis, possibly related to the degree of changes observed in the vestibulo-motor function they are implicated in. Indeed, whereas a majority of VO neurons remains tonic, in accordance with the limited changes reported in vestibulo-ocular circuits and reflexes known in amphibian species (for review see Straka and Dieringer, 2004), the expression of the two vestibular phenotypes was completely reversed in VS populations, from a majority of tonic 2°VN in larvae to a majority of phasic 2°VN in juvenile, to be associated with the complete remodeling of the vestibular control of posture that occurs during the metamorphosis (Olechowski-Bessaguet et al., 2020; Barrios et al., 2024). Thus, the next question was to identify by which mechanisms the expression of 2°VN phenotypes was differentially modified in distinct vestibular populations during that developmental period.

### Neurogenic origin of VS and VO neurons in juvenile

The first mechanism to be considered to explain the changes observed in the expression of 2°VN phenotypes was a neurogenic activity producing phasic and tonic neurons in different proportions between VS and VO populations from larvae to juvenile. To test this neurogenic hypothesis, BrdU pulse/chase experiments were first performed to identify the mitotic origin of vestibular neurons present at juvenile stage. Tadpoles were incubated 24h in BrdU at larval stages 42, 48, 50, 54, 58 or 65 (see experimental scheme in top Fig. 4A). After the pulse, animals were raised until post-metamorphosis stage 65 where either VS or VO neurons were retrogradely labeled in combination with both BrdU and Kv1.1 immunodetection (Fig. 4A). The total numbers of BrdU+ neurons (Fig. 4B_1_ and C_1_) and BrdU+/Kv1.1+ neurons (Fig. 4B_2_ and C_2_) were counted to determine the proportion of VS (Fig. 4B) and VO (Fig. 4C) neurons present at stage 65 and originating from either stages 42, 48, 50, 54, 58 or 65 (Fig. 4A-C). First, no VS or VO neurons were found BrdU+ when the pulse was performed at stage 58 (Fig. 4B_1_ and 4C_1_) or latter (not shown), demonstrating that no vestibular neurons in juvenile originated from a mitotic cell after stage 54. The latest mitotic origin of juvenile vestibular neurons was determined around stage 54, at early pro-metamorphosis, with only ∼5% of VS and ∼7% of VO neurons being BrdU+ (Fig. 4B_1_ and 4C_1_). Approximatively 10% of juvenile vestibular neurons originated from cells in division at stage 50 and about 20% from cells in division at stage 48 and stage 42 (Fig. 4B_1_ and 4C_1_).

**Figure 4.**
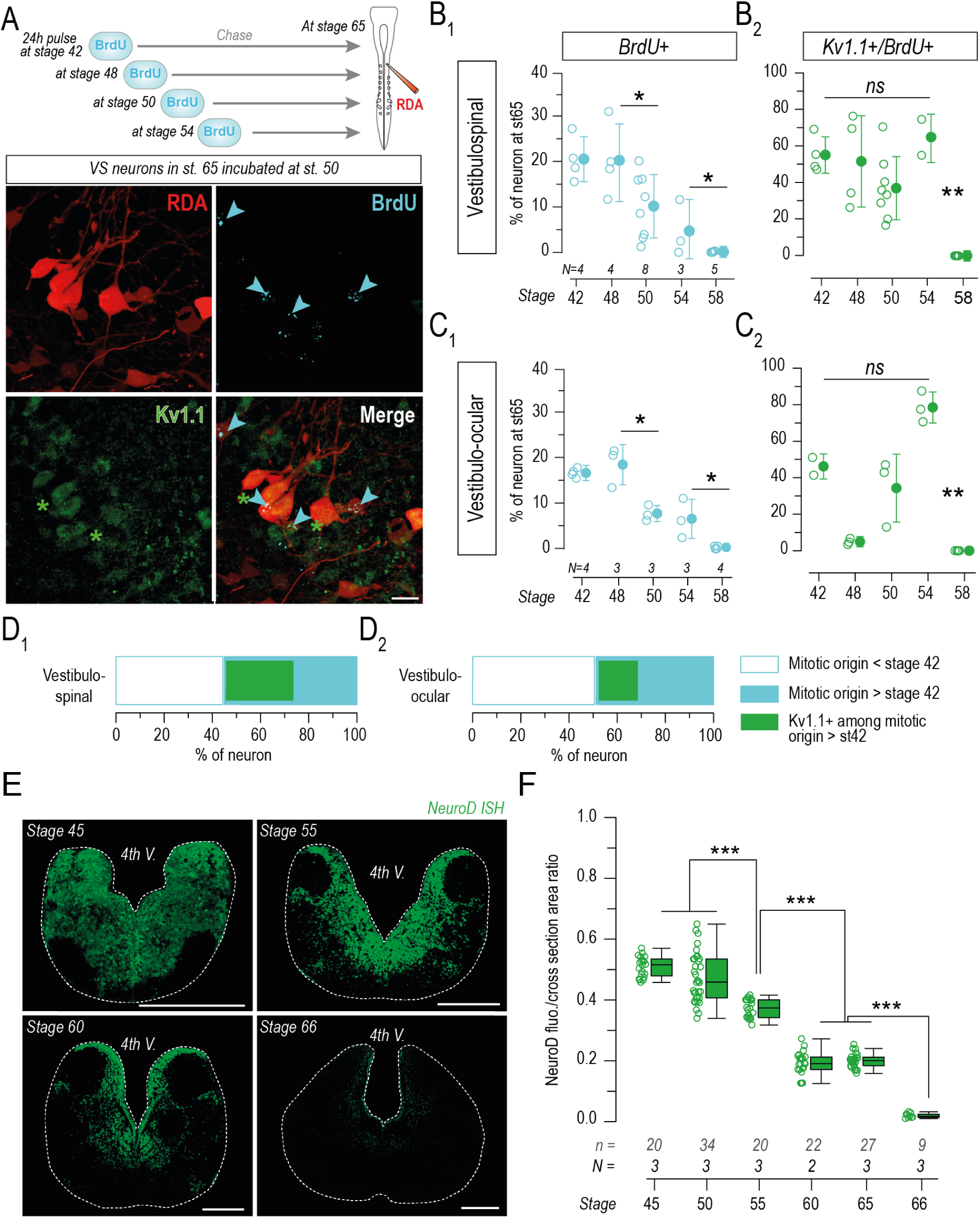
Neurogenic origin of juvenile vestibulospinal and vestibulo-ocular neurons. **A.** Schematic of the BrdU pulse/chase protocol to quantify the mitotic origin of vestibulospinal (VS) 2°VN in juvenile and example of confocal images (cross-sections) showing VS neurons retrogradely labeled with rhodamine dextran (RDA, in red) and immunodetection of Kv1.1 (green) and BrdU (blue) at juvenile stage 65 that were incubated with BrdU at larval stage 50. Blue arrowheads indicate BrdU positive VS neurons. Blue arrowheads with a green star indicate BrdU +/Kv1.1+ VS neurons. **B.** Averaged (±SD) proportions of BrdU positive (BrdU+; B_1_) and BrdU and Kv1.1 positive (Kv1.1+/BrdU+; B_2_) vestibulospinal neurons retrogradely labeled in juvenile from animal groups incubated in BrdU at larval stage 42, 48, 50, 54, 58. **C.** Averaged (±SD) proportions of BrdU+ (C_1_) and BrDU+/Kv1.1+ (C_2_) vestibulo-ocular neurons retrogradely labeled in juvenile from animal groups incubated in BrdU at larval stage 42, 48, 50, 54, 58. “N” in B-C indicates the number of animals. **D.** Cumulative proportions from B_1-2_ (D_1_) and from C_1-2_ (D_2_). Statistical tests: Percentages of BrdU+ (B_1_, C_1_) and Kv1.1+/BrdU+ (B_2_, C_2_) were compared with Khi² test (*=p<0.05, **=p<0.01). **E.** Confocal stack images of brainstem cross-sections showing cells labeled with the NeuroD *in situ* hybridization (ISH) at developmental stages 45, 55, 60, 66. **F.** Statistical area ratio of the NeuroD fluorescence in brainstem cross-sections at developmental stages 45, 50, 55, 60, 65 and 66. “n” indicates the total number of cross-sections used and “N” the number of animals for each group. Statistical tests: Assumption of normality was checked with Shapiro-Wilk test and sample variances compared with F-Test. NeuroD fluorescence areas were compared with ANOVA followed by Tuckey post-hoc test (***=p<0.005). “4th V.” = 4_th_ ventricle. Scale bar in A = 20µm and scale bar in E = 500µm.

This mitotic origin pattern was pretty similar between VS and VO neurons and was comparable with previous results (Van Mier et al., 1986). The proportion of Kv1.1+ neurons, presumably phasic, among BrdU+ VS neurons was relatively variable but did not display any significant evolution with the larval stage at which BrdU pulse was performed (Fig. 4B_2_). This proportion was even more variable in VO neurons over the different stages tested (Fig. 4C_2_). These data suggested that VS and VO neuronal populations in juvenile originated from neurogenesis profiles presenting different chronologies. Indeed, half of VS and VO neurons in juvenile (55.6% for VS neurons and 49.5% for VO neurons; the blue bar in Fig. 4D_1-2_) derived from mitotic cells between stage 42 and 54 whereas half of them originated from mitotic cells before stage 42 (Fig. 4D_1-2_). However, the proportion of Kv1.1+ neurons originating from larval stages 42-54 was higher in the VS population (∼50% of these neurons were Kv1.1+ representing 28% of the entire VS population in juvenile; green bar Fig. 4D_1_) than in the VO population (only ∼32% of these neurons were Kv1.1+ representing 16.3% of the entire VO population in juvenile; green bar Fig. 4D_2_). Consequently, it seems that the number of phasic, Kv1.1+, 2°VN in juvenile that derived from mitotic activity between stage 42 and 54 was more important in VS populations than in VO populations.

Conjointly to the mitotic origin, we also evaluated the global level of neuronal post-mitotic differentiation in the vestibular brainstem region by detecting the mRNA of the pro-neuronal D transcription factor (NeuroD; Fig. 4E; D’Amico et al., 2013) in order to precise the neurogenesis activity from stage 45 to stage 65. ISH labeling of the NeuroD marker revealed an important amount of neuronal post-mitotic progenitors in the brainstem at stage 45 and 50 (∼50% of the tissue; Fig. 4E-F). The NeuroD labeling decreased continuously from stage 55 to stage 66 where only a few NeuroD+ cells were detected in the brainstem (Fig. 4E-F), mostly in the vestibular region as previously reported by D’Amico and colleagues (2013). The ISH labeling of the NeuroD marker showed that most of the post- mitotic neuronal differentiation activity in the brainstem occurred until larval stage 55, concomitantly with the time course of the mitotic origin of vestibular neurons. Both results obtained from BrdU and NeuroD ISH labeling confirmed that the neurogenesis of VO and VS populations present in juvenile took place essentially before the pro-metamorphosis. However, despite the decrease of the NeuroD factor expression after stage 55, vestibular brainstem regions demonstrated a sustainable post-mitotic differentiation activity during the metamorphosis, potentially responsible for the changes in proportion of phasic and tonic phenotypes in VS and VO populations.

### Phenotype switch mechanism of tonic VS neurons into phasic VS neurons

Neurogenic activity is clearly involved in the change of proportion of tonic and phasic 2°VN during metamorphosis. However, this developmental mechanism cannot account for the entire process, suggesting that other mechanisms, like phenotype switch comparable to those one previously described in larval xenopus (Ernsberger and Spitzer, 1995; Dulcis et al., 2017; Lambert et al., 2018; Hammond-Weinberger et al., 2020) might operate conjointly. In this purpose, we used the computational neuronal modeling approach to evaluate the feasibility of a switch from a tonic neuron into a phasic neuron through a progressive up- and/or down-expression of membrane currents (Fig. 5 and 6). In a first step, we created a basic computational model for both tonic and phasic neuronal types with the currents known to be expressed in vestibular neurons, required to generate spikes and distributed differentially in distinct cellular compartments (Fig. 5A; Gittis et al., 2010). Second, because most of the VS tonic and phasic neurons were found to express a biphasic AHP (see Fig. 2A_3_-B_3_), BK and SK calcium-activated potassium currents, previously described to be responsible for biphasic AHP in vestibular neurons, were also included in the model (Fig. 5A-E and Sup. Fig. 5A; Smith et al., 2002; Iwasaki et al., 2008; Kolkman et al., 2011; Nelson et al., 2017). In addition to Kv1.1, the Kv7-M channel (Kv7.2/3 in the model) appeared to be a good candidate involved in the phasic phenotype (for review see Brown and Passmore, 2009). Its implication in fast adaptive firing patterns was reported in several neuronal types and several species (Bothe et al., 2024; Watanabe et al., 2017). Moreover, Kv7 and SK channels play a complementary function in the neuron excitability and synaptic excitation (Mateos-Aparico et al., 2014). Consequently, it appeared relevant to also consider the possible implication of the Kv7-related M current in the phasic VS phenotype. Importantly, we validate the presence of these different conductances in physiological vestibular neurons using pharmacological tests: BK and SK conductances with Iberiotoxin and Apamin (Sup. Fig. 5A-C), and Kv7.2/3 with XE991 (Fig. 5G-H). In addition, the presence of the Kv7.4 mRNA in central vestibular neurons was verified by performing an *in situ* hybridization labeling (Sup. Fig. 5D), demonstrating the combined presence of both Kv1.1 and Kv7-M currents at least in Kv1.1+ neurons.

**Figure 5.**
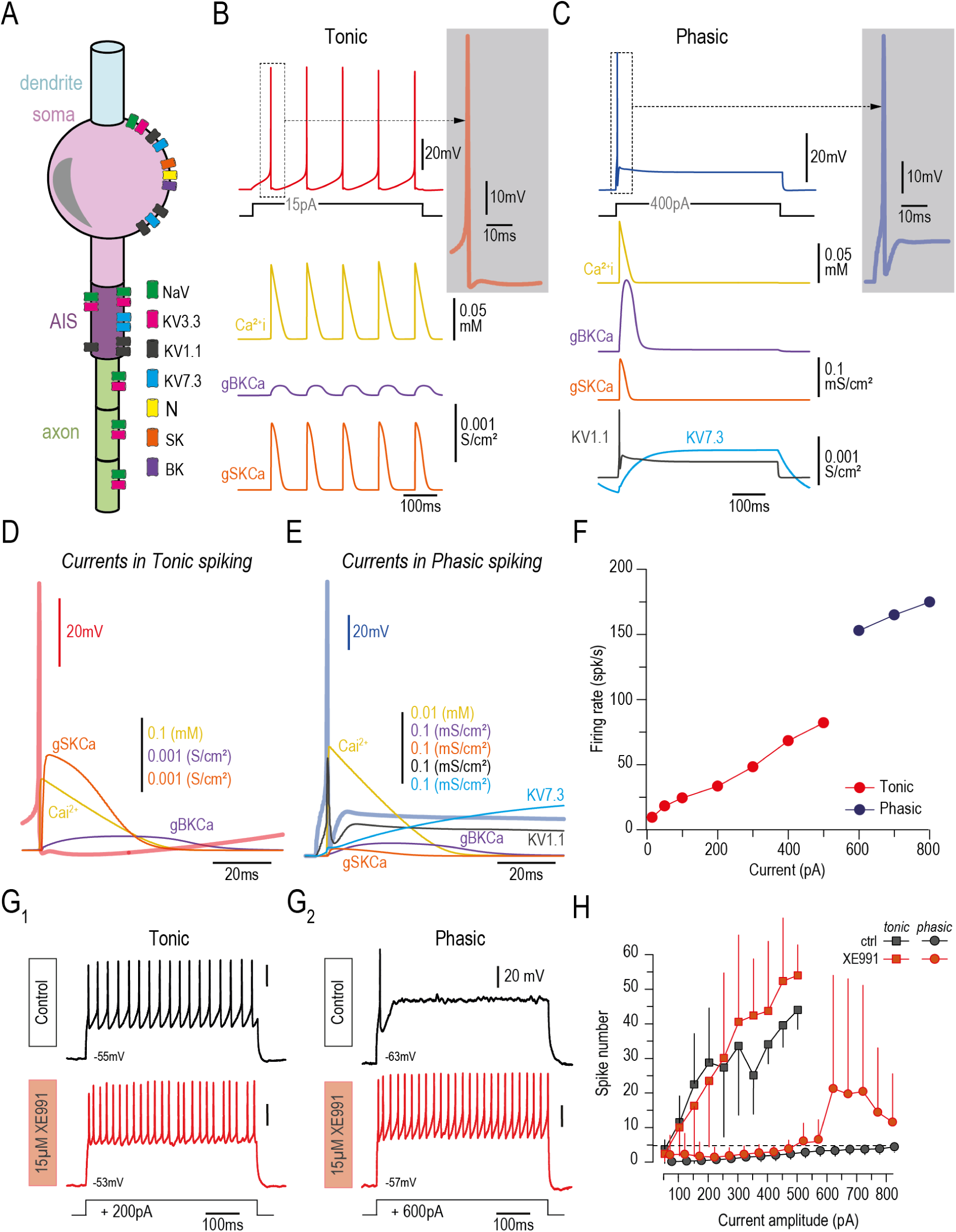
Neuronal architecture, currents distribution and density used to build tonic and phasic phenotype neuronal models. **A.** Schematic view of neuronal compartments with location of all currents used in 2°VN model. **B.** Low-induced current discharge of a tonic model with the variation of the intracellular Calcium (Ca²_+_i, in yellow) and the contributions of calcium-dependent BK and SK currents (gBKCa in purple and gSKca in orange). **C.** Low-induced current discharge of a phasic model with the variation of the intracellular Calcium (Ca²_+_i, in yellow) and the contributions of calcium-dependent BK and SK currents (gBKCa in purple and gSKCa in orange) and the voltage-dependent potassium currents Kv1.1 (in black) and Kv7.3 (in blue). **D-E.** Spike shape of a tonic neuronal model (D; red thick line) and phasic neuronal model (E; blue thick line) together with corresponding variation of intracellular calcium and contributions of BK, SK, Kv1.1 and Kv7.3 currents. **F.** Firing rate (in spike/s) of the tonic (red) and phasic (blue) 2°VN model in response to increasing current steps. **G.** Example of firing pattern of recorded tonic (G_1_) and phasic (G_2_) VS neurons before (control) and under bath application of XE991, blocking the Kv.7-M current. **H.** Averaged spike number in response to increasing current step injections before (ctrl) and during XE991 application in recorded tonic (squares) and phasic (dots) neurons. The horizontal dashed line represents the 5 spikes limit defining a phasic 2°VN neuron in amphibians (Straka et al., 2004; Beraneck et al., 2007).

**Figure 6.**
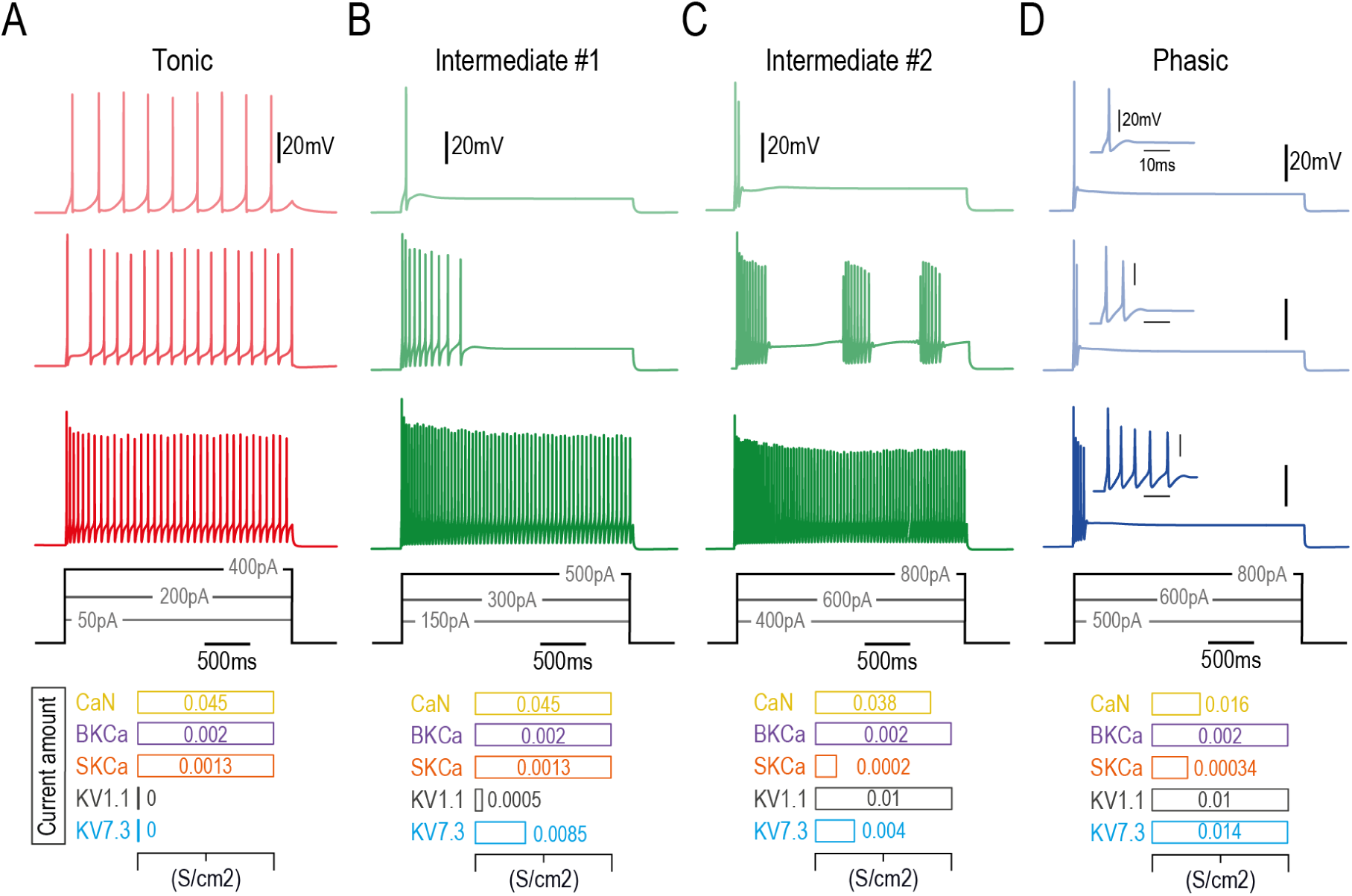
Switching from tonic to phasic neuronal model by progressively increasing Kv1.1 and Kv7.2/3 densities. A-D. Firing patterns at 3 different illustrative current steps when both Kv1.1 and Kv7.2/3 are null (A; in red), partially introduced (B-C; in green) and strongly expressed (D; in blue). The amount of individual currents used in each modeling configuration is indicated below traces in A-D.

By running simulations with our computational model, we were able to replicate the continuous firing pattern exhibited by real tonic VS neurons in response to current steps up to 500pA when Kv1.1 and Kv7.2/3 currents were absent and with a low quantity of SK current (Fig. 5B, D and F). Inversely, a higher amount of SK current together with the presence of both Kv1.1 and Kv7.2/3 induced a fast-adaptive discharge of 1-5 spikes similar to that expressed by real phasic VS neurons (Fig. 5C, E and F). Then we tested the possibility to turn a tonic VS neuron into a phasic neuron by changing the balance between the different key conductances involved (Fig. 6A-D). First, the introduction of a small amount of Kv1.1 and Kv7.2/3 affected the capacity of the tonic model to fire continuously in response to increasing current steps (Fig. 6A-B). In this modeling configuration, the neuron demonstrated an adaptive discharge until 400pA and was firing continuously only at 500pA (Fig. 6B).

Second, the same amount of Kv1.1 but with a reduced quantity of Kv7.2/3 induced a firing pattern comparable to that of phasic neurons until 500pA but showed an interrupted bursting activity at 600-700pA and a continuous firing only at 800pA (Fig. 6C). Thus, these simulations indicate that the progressive increase of Kv1.1 and Kv7.2/3 together with minimal adjustments of CaN and SK currents could allow the generation of firing patterns with intermediate features between tonic and phasic discharges. Furthermore, the model suggests also that the emergence of the phasic phenotype seems to be under the control of both Kv1.1 and Kv7.2/3. Indeed, increasing partly the amount of either Kv1.1 or Kv7.2/3 (Sup. Fig. 6A-B) only was not sufficient to reproduce intermediate discharge patterns comparable to intermediate model conditions #1 and #2 shown in figure 6. Altogether these different simulation results support the hypothesis that a switch from tonic to phasic discharge pattern can occur based on a change in the balance between certain conductances. Moreover, our modeling findings make the prediction that some VS neurons, when recorded during their transition period, could exhibit an “intermediate” discharge pattern no longer tonic anymore but not strictly phasic yet.

We directly tested this prediction by re-analyzing all patch clamp recordings performed on VS neurons. Interestingly, among a total number of 216 recorded neurons we identified 25 neurons exhibiting an intermediate discharge pattern, as illustrated in Figure 7 (Fig. 7A-C and Sup. Fig. 7A-C). This type of neurons was found exclusively in VS population, at all developmental stages examined and represented ∼10-13% of recorded neurons at each stage (Fig. 7D). Unlike tonic neurons, these neurons were not able to fire continuously in response to current step injections, at least for most of the step amplitudes tested (Fig. 7A-C). They rather presented an adapting discharge, but with a longer duration and a higher number of spikes than in phasic neurons (Fig. 7A-C and E). In addition, comparatively to phasic neurons, these intermediate neurons elicited a continuous firing activity during DTX-k or 4AP bath application (n=6; Fig. 7C_1-2_, sup. Fig. 7D), confirming the implication of the Kv1.1-related current in their adapting discharge pattern. We were also able to record one intermediate VS neuron under XE991 that fired continuously in the presence of the drug (Sup. Fig. 7E-F), confirming the potential implication of the Kv7-M current in this intermediate phenotype (as suggested by our model).

**Figure 7.**
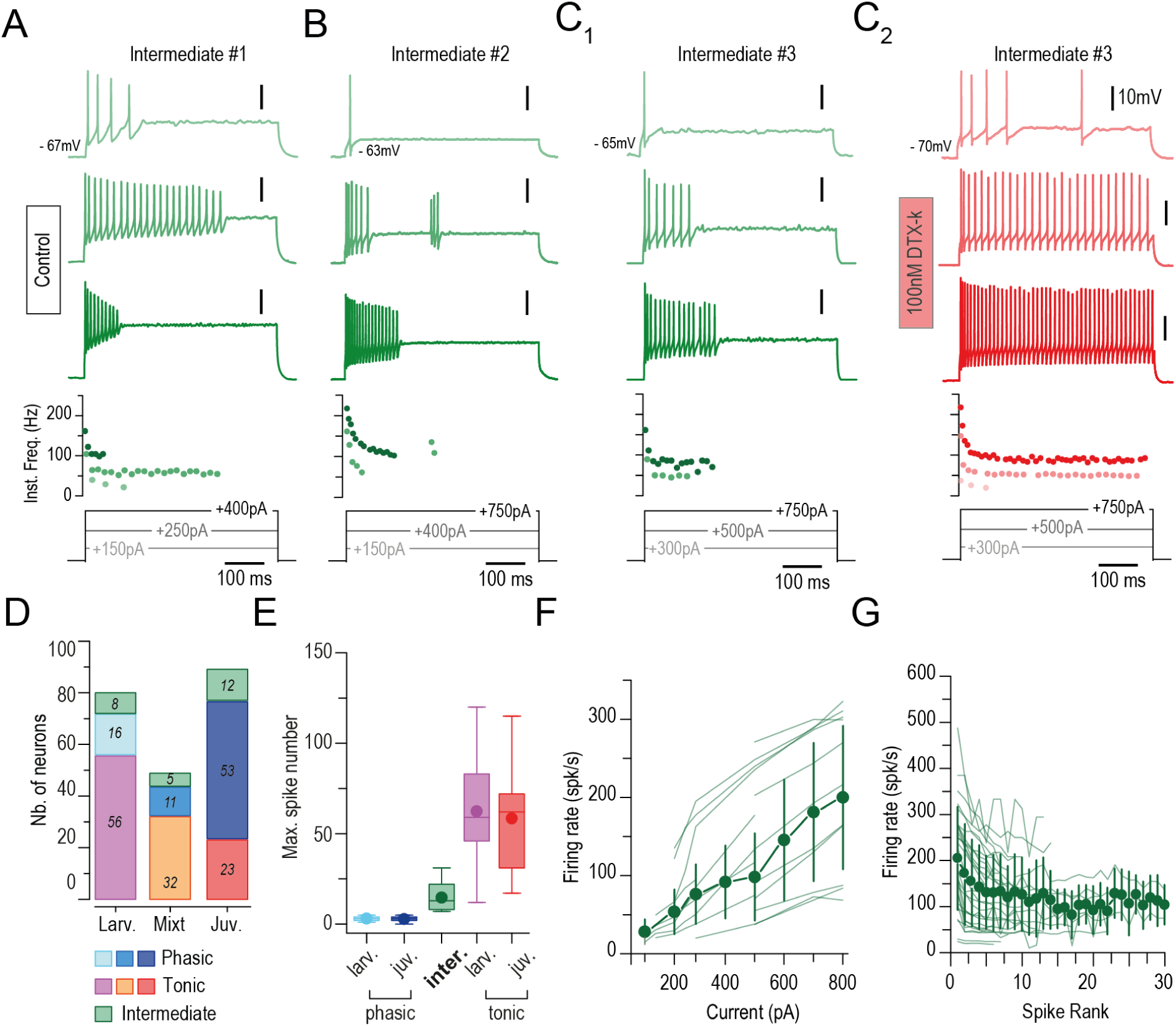
Vestibulospinal neurons with an intermediate firing pattern different from phasic and tonic phenotypes. A-C. Three examples of 2°VN presenting an intermediate firing pattern identical neither to the tonic phenotype nor to the phasic one. These neurons became tonic under bath application of DTX-k or 4AP as shown for the neuron presented in C_1_ (control) and C_2_ (DTX). **D.** Number of phasic, tonic and intermediate phenotypes found in VS neurons at larval, mixt (climax) and juvenile stages. **E.** Averaged maximum spike numbers elicited by phasic, intermediate and tonic VS neurons at larval and juvenile stages. Larval and juvenile intermediate neurons were pooled due to the small number of neurons and the absence of significant difference in maximum spike number. **F-G.** Individual and averaged firing rate (±SD; in spike/s) plotted against the injected current step (F) or against the spike rank (the position of the spike in the spike train, G).

Even if we did not record a lot of neurons exhibiting this intermediate phenotype, they seem to represent a distinct neuronal group found during all the metamorphosis period and obviously specific to vestibular neurons projecting to the spinal cord. In contrast with tonic and phasic neurons, these intermediate neurons presented an important heterogeneity in their discharge features (Fig. 7A-C and 7F-G) and this independently of the developmental stage (Sup. Fig. 7A-C). Such a discharge heterogeneity certainly relies on a more variable amount of some currents (including the Kv1.1 and/or Kv7 currents) in intermediate neurons than in either tonic or phasic neurons. Therefore, the sustainable presence of this intermediate phenotype in VS neurons argues for a possible functional switch mechanism turning tonic neurons into phasic neurons from larval to juvenile stage, based on down- or up-expression of ion channels progressively changing the continuous non-adapting discharge of tonic neurons into the fast-adapting phasic pattern through a transitory discharge profile.

To further identify potential mechanisms underlying this developmental switch in membrane properties of VS neurons we investigated the possibility for tonic neurons to progressively synthetize functional Kv1.1 channel from an internal mRNA stock in order to acquire features of phasic neurons.

For that we first tested whether some VS neurons projecting to the spinal cord in juvenile were already present at larval stages. To do so, dextran dye crystals were injected unilaterally in the spinal cord at larval stage 53, around spinal segments 3 to 6, with a limited invasive surgery compatible with the animal survival (Fig. 8A). After metamorphosis, an additional retrograde labeling of VS neurons was performed from the rostral spinal cord, combined with a Kv1.1 immuno-labeling (Fig. 8A). We found that 15±6.9% of juvenile VS neurons were also labeled with the dextran dye injected at larval stage, indicating that these neurons were already differentiated and projecting to the spinal cord before metamorphosis (Fig. 8B). The great majority of these double labeled VS neurons were Kv1.1+ (71.3±20.8%; Fig. 8B), supporting the possibility that these neurons could indeed express a phasic phenotype at juvenile stage. Based on 1/ the low proportion of phasic VS neurons found in larvae (∼20%; see Fig. 2) and 2/ the non-subtype targeted labeling of VS neurons at larval stage, the probability that all VS neurons double labeled in juvenile exhibited already the phasic phenotype in larvae appears to be improbable. Consequently, we presumed that some of these phasic (Kv1.1+) juvenile VS neurons projecting to the spinal cord before metamorphosis did not express Kv1.1 at larval stage, and would consequently exhibit a tonic discharge.

**Figure 8.**
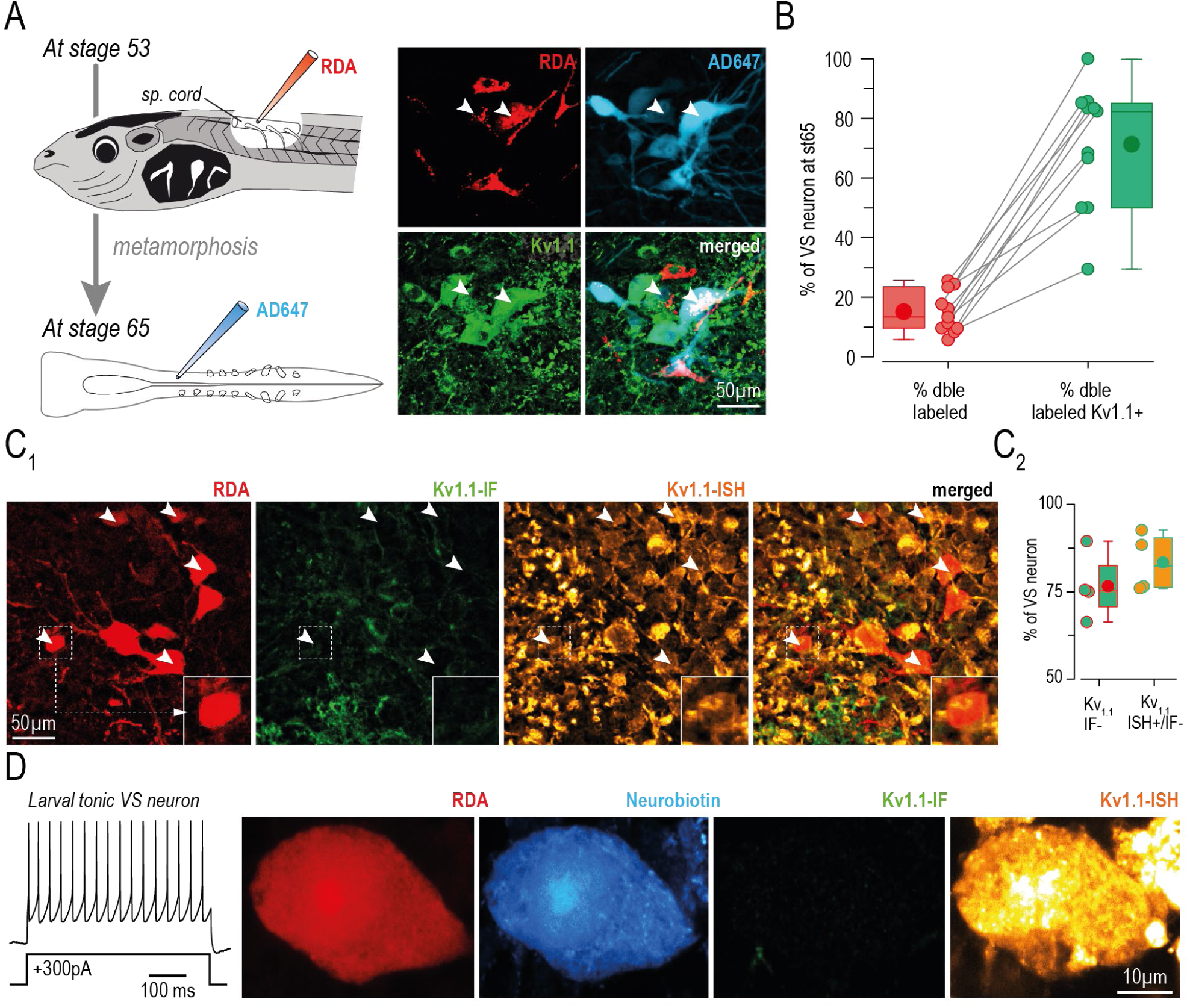
Larval tonic vestibulospinal neurons express the Kv1.1 mRNA but not the Kv1.1 protein. **A.** Schematic view of the double neuronal tracing method to track juvenile (stage 65) vestibulospinal (VS) neurons already projecting in the spinal (sp.) cord at larval stage (stage 53) and example of confocal image of VS neurons in juvenile retrogradely labeled in larvae with Rhodamine dextran amine (RDA, red) dye and with Alexa 647 dextran amine (AD647, blue) dye at stage 65, either Kv1.1 positive (Kv1.1+, green, indicated by white arrowhead) or negative (Kv1.1-). **B.** Proportion of double (dble, red circles) labeled VS neurons, corresponding to juvenile VS neurons already present in larvae, and proportion of Kv1.1+ neurons among these double labeled neurons (green circles). **C.** Example of confocal image (C_1_) and proportion (C_2_) of VS neurons retrogradely labeled with RDA, not labeled with immuno-fluorescent Kv1.1 (Kv1.1-IF-) but labeled with *in situ* hybridization against the Kv1.1 mRNA (Kv1.1-ISH+; indicated by white arrowheads). **D.** Tonic VS neurons recorded in response to current step at larval stage (trace on the left), retrogradely labeled with RDA, backfilled intracellularly with Neurobiotin during the path-clamp recording without immuno-fluorescent labeling (green) of Kv1.1 protein but labeled with ISH labeling for Kv1.1 mRNA (orange).

In a more demonstrative way, we searched for the presence of the Kv1.1 related mRNA in larval tonic VS neurons despite the absence of a functional Kv1.1 channel. To do so, Kv1.1 immunofluorescence and *in situ* hybridization were combined on VS neurons retrogradely labeled at larval stage (Fig. 8C_1_). As already shown in Figure 3, most of larval VS neurons were Kv1.1 immune-negative (76.6±9.5%; Fig. 8C_2_), demonstrating the absence of Kv1.1 channel expression in these cells. However, *in situ* hybridization revealed that most of these neurons were positive for Kv1.1 mRNA (Fig. 8C_2_). Even more strikingly, we performed the same combined Kv1.1 immunofluorescence and *in situ* labeling on larval neurons physiologically identified with patch-clamp recording (Fig. 8D). Again, we found some larval VS neurons (n=5), showing a tonic discharge pattern, identified with the neurobiotin intracellular labeling, negative for the Kv1.1 immunofluorescence signal but positive for the *in situ* Kv1.1 labeling (Fig. 8D). All together, these findings unambiguously demonstrated that larval tonic VS neurons expressed the mRNA of the Kv1.1 channel but not the mature Kv1.1 channel protein. The presence of the Kv1.1 channel protein at later developmental stages will thereafter sustain the expression of the typical phasic phenotype, thus participating to the maturational process of the vestibular neurons discharge properties.

In conclusion, our computational approach demonstrated that a small amount of changes in membrane currents balance could be sufficient to turn the discharge pattern from tonic to phasic. These modeling results (shown in figures 5 and 6) in combination with the data obtained on physiological VS neurons (figures 7 and 8) are strong and convincing arguments for the hypothesis of a phenotype switch mechanism transforming tonic neurons into phasic ones. Such a developmental mechanism might participate, conjointly with the neurogenesis, to reverse the proportion of tonic and phasic specifically in the VS neuronal population during frog metamorphosis.

## DISCUSSION

Our findings reveal that VS and VO 2°VN present different maturation profiles during frog metamorphosis in direct correlation with the degree of adaption in vestibular-driven motor ocular and postural responses imposed by functional changes occurring during this developmental period. Indeed, discharge properties of VO 2°VN is highly comparable in both larvae and froglets, with a large majority of neurons exhibiting a tonic discharge pattern, concomitantly with the mostly unchanged vestibular-ocular reflexes. Inversely, VS 2°VN present more pronounced developmental changes from larval to juvenile stages. First, the global firing activity of tonic VS neurons is lower in juvenile. Second and more importantly, the respective proportions of phasic and tonic phenotypes in the VS population are completely reversed, switching from a majority of tonic neurons in larvae to a majority of phasic neurons in juvenile. These important changes in VS 2°VN are to be associated to the complete transformation of the posturo-locomotor system during the metamorphosis, where the limb-based thoraco-lumbar system replaces progressively the tail-based axial system, at both neural and musculoskeletal levels. These differential maturation profiles seem to rely partly on the neurogenic activity in vestibular neuronal subpopulations but is supported also by a conjoint post-differentiation phenotype switch mechanism, at least in VS neurons, turning some larval tonic 2°VN into phasic neurons in juvenile.

### Maturation of 2°VN subpopulations reflects the different degree of reorganization in vestibular postural and oculomotor responses during metamorphosis

Motor control and sensory-motor transformation tasks depend, at least partly, on discharge dynamics expressed by the neuronal populations involved, specifically tuned to produce the appropriate behavioral output. Such a cellular-behavioral adequation was already highlighted in various motor systems, like in spinal microcircuits responsible for swimming in zebrafish (Ampatzis et al., 2014; Björnfors and El Manira, 2016; for review see Picton et al., 2025), or the vestibulo-ocular pathways in amphibian (Straka et al., 2009, Beraneck and Straka, 2011). The maturation profiles in 2°VN populations described in the present study fully endorses the changes encountered in the posturo-locomotor and the oculomotor behaviors and the adaptation of the vestibular control they both involve. In tadpoles, postural control consists in a slow but regular tail undulation to maintain the body in the water column. In contrast, froglets alternate long motionless phases on the ground with brief and fast asymmetrical movements (Cheung et al., 2005 and 2009; Beyeler et al., 2013; Reilly et al., 2015; Li et al., 2021). These two postural strategies rely on completely different vestibulospinal motor responses that produce either a constant low frequency swimming activity in larvae and rapid, compensatory, reflexive movements of both trunk and limbs in adults (Ehrlich and Schoppik, 2019; Olechowski-Bessaguet et al., 2020; Barrios et al., 2024). In this context, tonic VS neurons, with a noticeable spontaneous activity for 25% of them, appear to adequately sustain a vestibular motor command capable of maintain a slow rhythmic swimming activity, to constantly insure the body orientation and position in the water. In contrast, phasic VS 2°VN together with a minimum proportion of tonic neurons exhibiting a low firing rate and no spontaneous activity would be beneficial for a more adaptive postural activity, based on a flexible and complex coordination of trunk and hindlimbs, like observed in froglet. Strikingly, 80% of VS neurons in adult land frog also exhibit the phasic phenotype (Straka et al., 2004). Therefore, the important changes observed in the VS neuronal population accompany the complete reorganization of the spinal motor network and supports the radical change of posturo-locomotor behavior during the frog metamorphosis.

In contrast with the posturo-locomotor system, eye muscles and extraocular motor nuclei are not deeply reorganized during metamorphosis and visual and vestibular gaze-stabilizing reflexes are highly similar from larval to adult stages (Dieringer and Precht, 1982; Horn et al., 1986; Straka and Dieringer, 1991; Straka et al., 1998; Lambert et al., 2008; Dietrich et al., 2017; Gravot et al., 2017; Bacqué-Cazenave et al., 2022). Hence, frogs achieve a VOR composed mainly of smooth compensatory eye movements counteracting slow head position changes and little of fast head velocity adaptation. As a consequence, exhibiting a vestibulo-ocular population dominated by tonic neurons, acting as low-pass filter position signal encoders (Beraneck et al., 2007; Straka et al., 2009), would sustain a “slow dynamic” VOR at both larval and juvenile stages. Therefore, in contrast to the VS population, it appears relevant to find mostly tonic phenotype in the VO neuronal group without noticeable changes in electrophysiological properties before and after metamorphosis. Our results are based on the ipsilateral oculomotor-projecting VO group, implicated in VOR components very well conserved from larvae to froglet (Horn et al., 1986). However, we cannot exclude that other VO subpopulations mature differently, like VO neurons projecting on Abducens nucleus and involved in the horizontal angular VOR, a VOR modality with a developmental pattern different than the otolith VOR. Also, phasic and tonic proportions in VO neuronal groups could be attributed to inter-species differences. Indeed, hunting strategy in terrestrial frog, requiring more fast eye movements, could justify a higher proportion of phasic VO neurons than in a purely aquatic species like the Xenopus toad. Even if it remains hypothetical, this last explanation would also show a specific adaptation, not only during the development but also through the amphibian evolution, of 2°VN phenotypes according to specific needs of vestibulo-motor reflexes.

### Developmental mechanisms responsible for phenotype proportion changes in 2°VN

Previous studies investigated neurogenic activity in vestibular domains during larval life. For instance, some of these past reports showed that 1/ all 2°VN projecting to the spinal cord in froglet originated from a larval period between stage 10 and 50 (Van Mier et al., 1986) and 2/ neuronal differentiation activity, quantified by NeuroD ISH, was minimal in the brainstem of juvenile Xenopus and preferentially located in the vestibular area (D’Amico et al., 2013). The present study confirms these results for VS neurons but also VO neurons, suggesting a similar neurogenic temporal profile where most of the neurogenic activity occurs until stage 54. The pool of progenitors present at this pro-metamorphosis stage will enter in a post-mitotic differentiation process specific to each 2°VN subpopulations. In this context, neurogenic activity alone could be sufficient to insure the small increased of phasic VO neurons observed in juvenile. However, even if we reported a higher proportion of Kv1.1+ neurons (presumably phasic) deriving from mitotic cells between stage 42 and 50 in VS compared to VO populations (Fig. 4), surprisingly, it was not sufficient by itself to explain the complete reversion of proportion of tonic and phasic 2°VN in the VS population. Therefore, an alternative developmental mechanism might also be involved in the massive increase of phasic VS neurons during metamorphosis. Developmental-related neuronal switch mechanisms have been previously reported in Xenopus, but mostly for neurotransmitter phenotype changes (Dulcis et al., 2017; Spitzer, 2017; Lambert et al., 2018). In addition, recycling of larval neurons in adult stages have been also described, at least at the motoneuronal level (Barnes and Alley, 1983; Alley and Barnes, 1983; Herrera et al., 1991; Omerza and Alley, 1992; Alley and Omerza, 1998). Consequently, a cellular mechanism that transforms larval tonic neurons into phasic neurons, by up- and/or down regulating the expression of key ions channels appeared to be a good additional source to increase the number of VS phasic neurons during frog metamorphosis. Supporting this idea, previous observations reported maturation of brainstem neurons by regulating the expression of Kv1.1 or Kv7-M currents involved in adaptive firing patterns. Indeed, Watanabe et al (2017) showed that Mauthner cell discharge dynamic becomes more phasic during the early larval fish development, with a bursting discharge (even tonic for some stimulation values) at 2dpf and the firing of a single spike at 4-7dpf. Acquisition of this highly phasic pattern, comparable to phasic 2°VN in xenopus froglet, is due to a conjoint increase of both Kv1.1 and Kv7.4. Additionally, a change in electrophysiological phenotype in tangential vestibular neurons has been previously reported during the chick perinatal period where 2°VN exhibited a phasic discharge dynamic, expressing Kv1.1 channel in embryo but a tonic phenotype without kv1.1 after hatchling (Popratiloff et al., 2003). More recently, even if it was not in a developmental context, a functional switch has been reported in spinal motoneurons of rattlesnake by activation/blockade of the Kv7.2/3 channel (Bothe et al., 2024). Motoneurons involved in the rattling behavior exhibited an adaptive firing activity, similar to phasic 2°VN, whereas motoneurons responsible for locomotion presented a continuous discharge pattern, like in tonic 2°VN. Interestingly, the blockade of the Kv7.2/3 turned the rattling motoneuronal discharge from an adaptive to a non-adaptive firing pattern, similar to that expressed in locomotor motoneurons. Inversely the enhancement of the Kv7.2/3 provides the reverse effect on the locomotor motoneuron firing rate.

Therefore, the presence of Kv1.1 and Kv7-M currents in the neuronal adaptative firing pattern of central vestibular neurons demonstrated in the present study fits perfectly with all this previous literature and confirmed that this current implication might be commonly used to produce such a discharge dynamic in the central nervous system. In addition, the implication of the Kv1.1 and Kv7-M currents to trigger adaptive-firing in brainstem neurons could be an evolutionary conserved feature. Indeed, this combination seems to be present in several vertebrates’ lineages as suggested by the close phylogenetic link between fish and amphibian for Kv7.2, Kv7.3 and Kv7.4 genes revealed by Wanatabe and colleagues (2017). Our data in xenopus reinforce this hypothesis by bringing another example of neuronal population with the same molecular characteristic, at least in the brainstem. More importantly, our results strongly suggest the existence of a developmentally-induced switch that do not change the neurotransmitter switch of the phenotype, as described until now (Dulcis et al., 2017; Spitzer, 2017; Lambert et al., 2018), but rather modify the firing pattern involving specific membrane properties and consequently the computational and integrative capacity of the neuron. This would constitute a novel mechanism to consider in questioning about the neuronal plasticity.

Altogether, our findings brought a new insight in the understanding of the maturation and the adaptive plasticity of the vestibular system during the frog metamorphosis. We showed that VS and VO 2°VN did not follow the same maturation profile from larval to juvenile stage, highly related to the specific and different changes occurring in the posturo-locomotor and oculomotor systems behaviors. These specific maturation profiles seem to be determined by different developmental processes and, therefore, push to investigate electrophysiological phenotypes in first born 2°VN at early larval stages. More broadly, this study proposes working hypothesis to investigate neuronal networks reorganization in crucial functions like locomotion, respiration or vocalization occurring during metamorphosis, a developmental strategy common to amphibians, insects and mollusks or to comparable maturational periods with an important remodeling.

## Acknowledgements

This work was supported by the Centre National de la Recherche Scientifique, and grants from the Agence Nationale pour la Recherche (*ANR-22-CE37-0002-01 LOCOGATE* and *ANR-22-CE16-0004-02 MOTOC*, F.M. Lambert), the Fondation pour la Recherche Médicale (*Equipe FRM 2017-2022*, M. Thoby-Brisson, DEQ20170336764). Authors are grateful to Gladys Alfama (PhyMA) for probe synthesis, to Anne-Laure Gaillard (PhyMA) for teaching L. Cardoit the FISH technique, to Lionel Parra-Iglesias (INCIA) for taking care of the animals.

---

**Supplemental figure 2.**
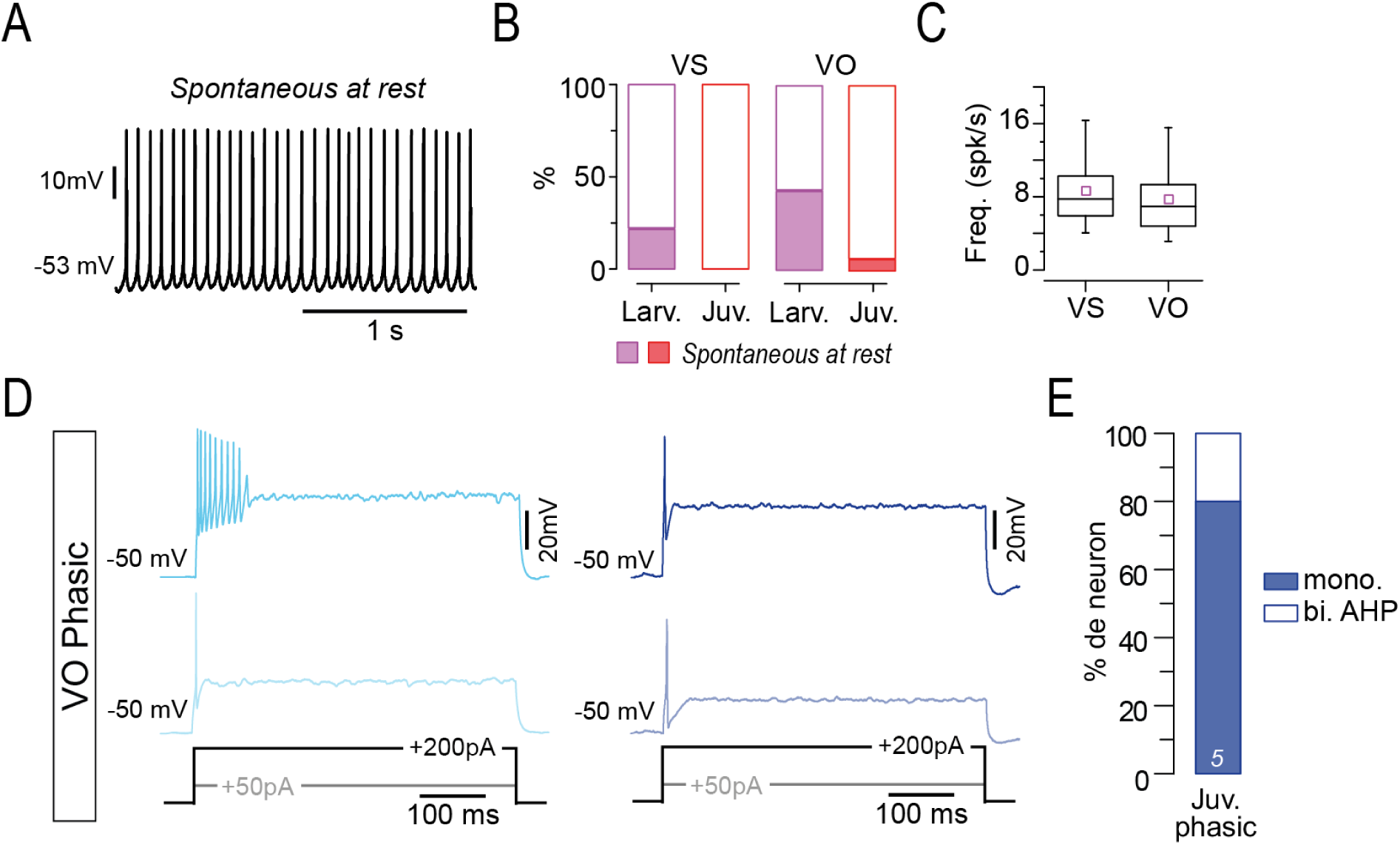
**A.** Examples of spontaneous firing activity at rest in larval tonic VS neurons. **B.** Percentage of tonic vestibulospinal (VS) and vestibulo-ocular (VO) neurons presenting a spontaneous firing activity at rest in larvae (Larv., purple) and juvenile (Juv., red). **C.** Averaged instantaneous (inst.) and mean firing rate (in spike/s, spk/s) of spontaneous firing activity at rest in larval tonic VS and VO neurons. **D.** Examples of discharge dynamics in larval (left, light blue) and juvenile (right, blue) phasic VO neurons in response to increasing positive current step injections during patch-clamp recording. **E**. Proportion of mono- and biphasic AHP for phasic VO neurons in juvenile.

**Supplemental figure 3.**
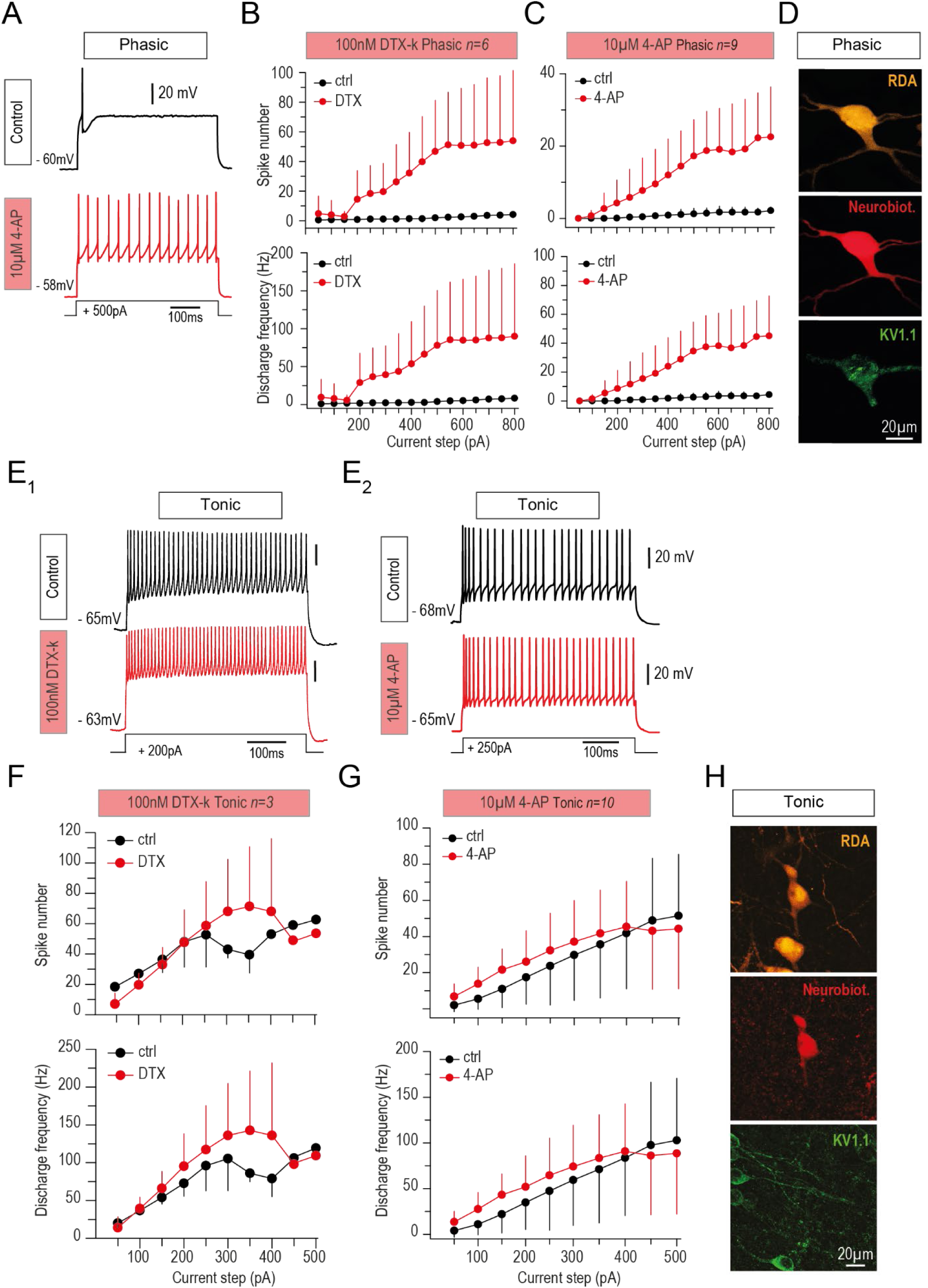
**A.** Firing pattern of phasic 2°VN during control (top black trace) and 10µM 4-AP bath application (bottom red trace). **B-C.** Averaged spike number (top graphs) and firing rate (bottom graphs, in spike/s, spk/s) of phasic 2°VN during control and DTX-k (B) or 4-AP (C) application. **D.** Confocal images of phasic 2°VN labeled with retrograde injection of rhodamine dextran amine tracer (RDA, top picture; VS neurons shown in example), with streptavidin-revealed neurobiotin injected through the recording electrode (red, middle picture) and with Kv1.1 immunolabeling (green, bottom picture). **E.** Firing pattern of tonic 2°VN during control (top black traces) and DTX-k (E_1_) or 4-AP (E_2_) bath application (bottom red traces). **F-G.** Averaged spike number (top graphs) and firing rate (bottom graphs, in spike/s, spk/s) of tonic 2°VN during control and DTX-k (F) or 4-AP (G) application. **H.** Confocal images (scale bar = 20µm) of tonic 2°VN labeled with retrograde tracer injection of RDA (RDA, top picture VS neuron shown in example), with streptavidin-revealed neurobiotin injected through the recording electrode (red, middle picture) and with Kv1.1 immunolabeling (green, bottom picture).

**Supplemental figure 5.**
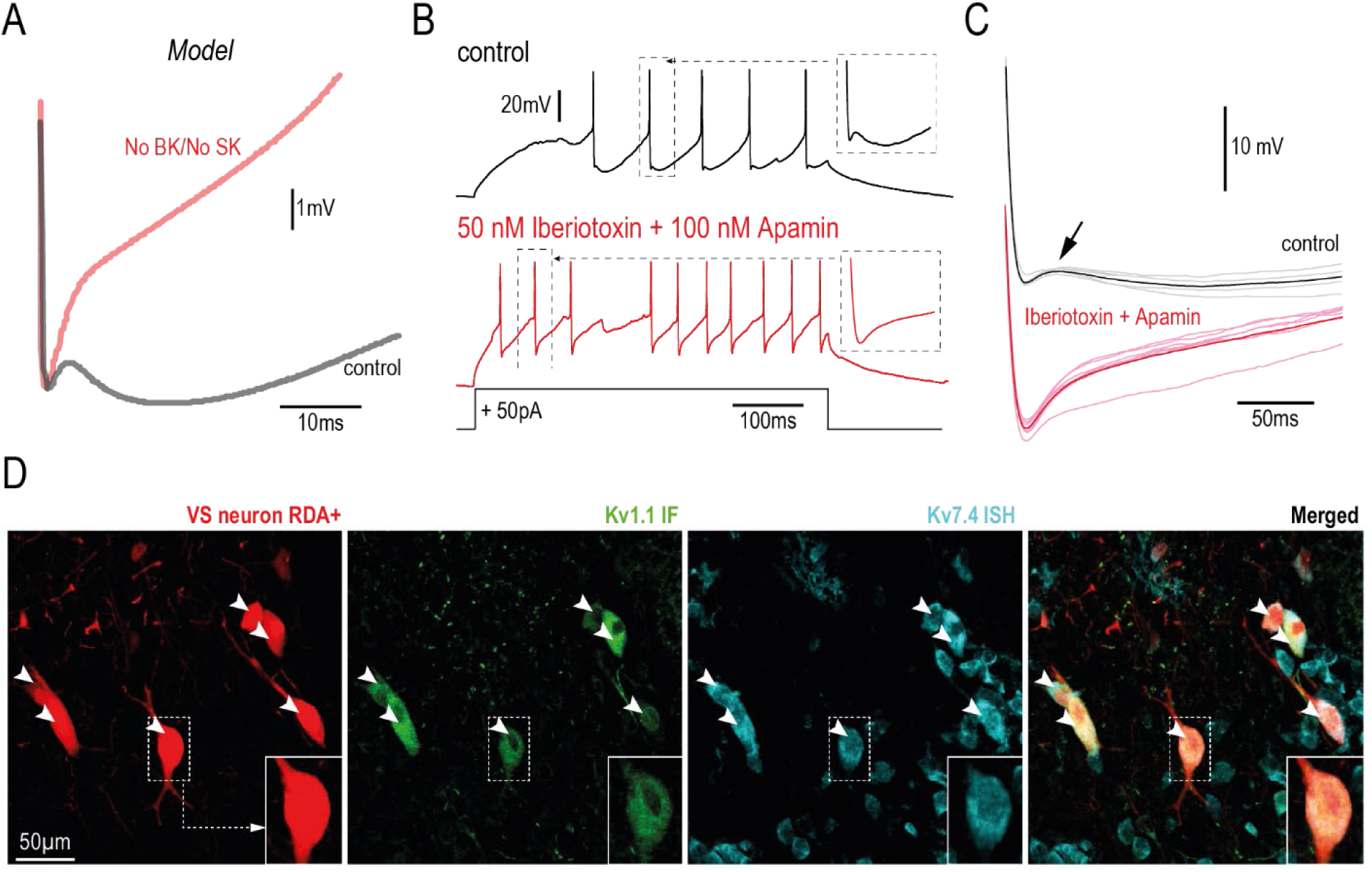
**A.** Biphasic after hyperpolarization (AHP) spike shape generated by the NEURON simulation of a 2°VN model in presence (black) or absence (red) of BK and SK currents. **B.** Firing pattern of a recorded tonic VS neurons in control (black upper trace) and during bath application of Iberiotoxin and Apamin (red bottom trace). **C.** Individual (thin lines) and averaged (thick line) shape of the spike after-hyperpolarization phase from the neuron shown in B. The black arrow indicates the biphasic AHP visible in control that disappears during combined Iberiotoxin and Apamin application. **D.** Example of confocal image of VS neurons retrogradely labeled with RDA in juvenile, labeled with immuno-fluorescent Kv1.1 (Kv1.1-IF) and labeled with *in situ* hybridization against the Kv7.4 mRNA (Kv7.4-ISH; indicated by white arrowheads).

**Supplemental figure 6.**
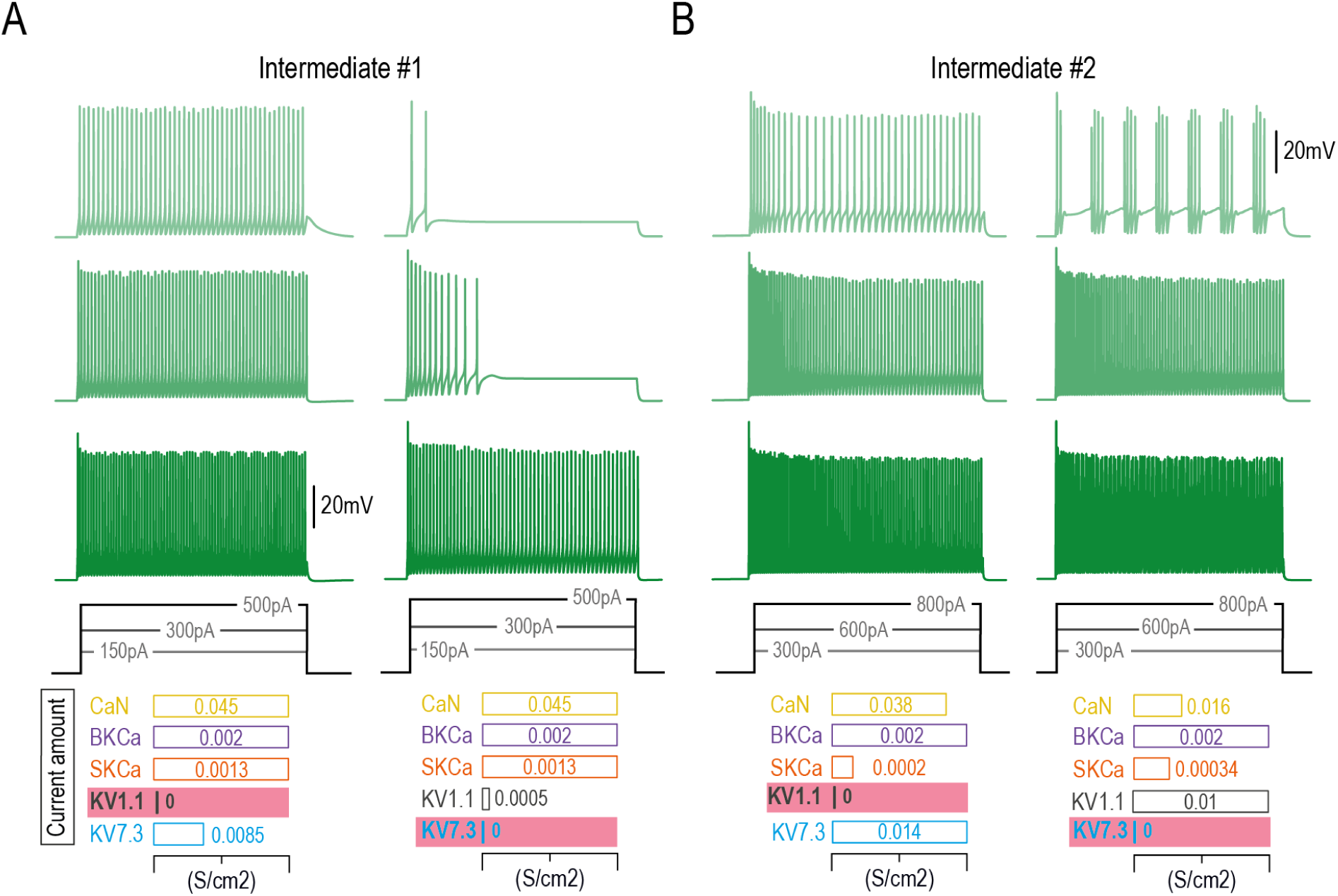
Firing patterns of modeling configurations presented in Figure 5B (A) and 5C (B), at 3 different illustrative current steps, when either Kv1.1 or Kv7.2/3 were maintained null (highlighted in red).

**Supplemental figure 7.**
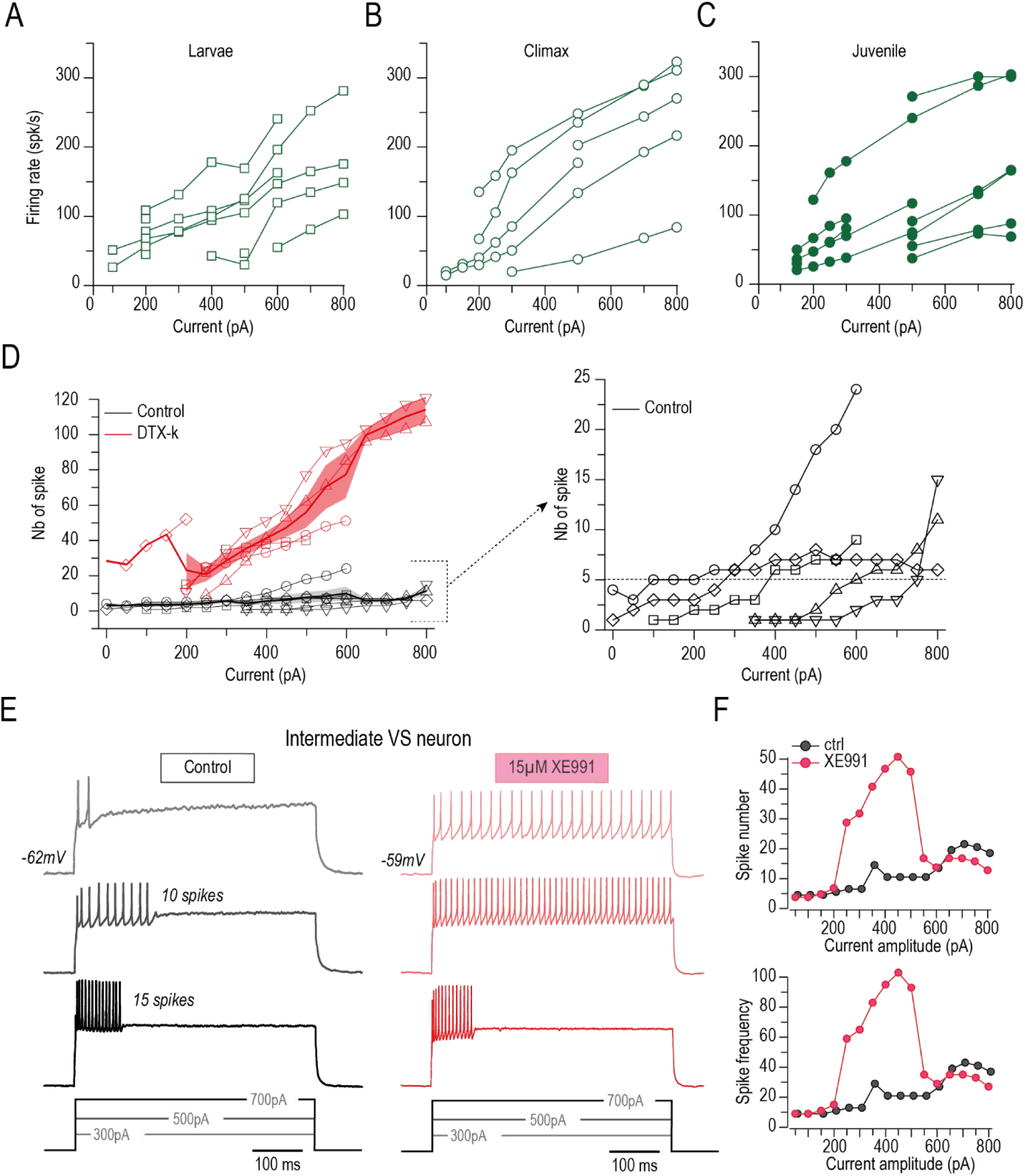
**A-C.** Firing pattern in response to increasing current step injections from individual intermediate VS neurons found in larvae (A), climax (B) and juvenile (C) stages. **D.** Number (Nb) of spike from individual intermediate VS neurons and average Nb of spike (+/- SEM) in control (black) and during DTX-k (red) bath application. The right plot in D is a scaled view of the control values shown in the left plot. The horizontal dashed line represents the 5 spikes limit corresponding to phasic phenotype. **E.** Firing pattern in response to increasing current step injection from one individual intermediate VS neuron in control (black) and during XE991 (red) bath application. **F.** Spike number and frequency in response to increasing current step injections of the neuron shown in E.

